# Shaping human gut community assembly and butyrate production by controlling the arginine dihydrolase pathway

**DOI:** 10.1101/2023.01.10.523442

**Authors:** Yiyi Liu, Yu-Yu Cheng, Jaron Thompson, Zhichao Zhou, Eugenio I. Vivas, Matthew F. Warren, Federico E. Rey, Karthik Anantharaman, Ophelia S. Venturelli

## Abstract

The arginine dihydrolase pathway (*arc* operon) present in a subset of diverse human gut species enables arginine catabolism. This specialized metabolic pathway can alter environmental pH and nitrogen availability, which in turn could shape gut microbiota inter-species interactions. By exploiting synthetic control of gene expression, we investigated the role of the *arc* operon in probiotic *Escherichia coli* Nissle 1917 on human gut community assembly and health-relevant metabolite profiles *in vitro* and in the murine gut. By stabilizing environmental pH, the *arc* operon reduced variability in community composition across different initial pH perturbations. The abundance of butyrate producing bacteria were altered in response to *arc* operon activity and butyrate production was enhanced in a physiologically relevant pH range. While the presence of the *arc* operon altered community dynamics, it did not impact production of short chain fatty acids. Dynamic computational modeling of pH-mediated interactions reveals the quantitative contribution of this mechanism to community assembly. In sum, our framework to quantify the contribution of molecular pathways and mechanism modalities on microbial community dynamics and functions could be applied more broadly.

## INTRODUCTION

The human gut microbiome substantially expands our genome’s capabilities by encoding a myriad of unique metabolic pathways^1–3^. Certain metabolic pathways are frequently found in constituent members of gut microbiota, such as the enzymes to produce acetate^4^. However, other specialized metabolic pathways such as production of butyrate and trimethylamine are present in a subset of gut species^5,6^. A given specialized metabolic pathway can be harbored by phylogenetically distant species and can play key roles in human health and disease^7^. In addition to directly influencing the host, these pathways can shape the dynamics and functions of gut microbiota. Understanding the quantitative contributions of these pathways on community dynamics and functions could reveal new influential molecular control knobs for shaping community states.

The arginine dihydrolase pathway (encoded by the *arc* operon or arginine deiminase system) is a specialized metabolic pathway that converts arginine into several products including ammonia, which could impact the gut community^8^. The *arc* operon has been shown to influence pH homeostasis in oral biofilms^9^ and its prevalence has been studied in oral species^10^, strains of *Escherichia coli*^11^ and *Eggerthella lenta* (*E. lenta*)^12^. Specifically in the gut, this pathway is harbored by diverse human gut species enabling utilization of arginine as an energy source under aerobic and anaerobic conditions^13–15^. The *arc* operon in *Enterococcus faecium* enhanced *Clostridioides difficile* toxin production through arginine conversion and ornithine cross-feeding^16^. By releasing ammonia, the *arc* operon can increase environmental pH and provide a nitrogen source that could benefit the community in nitrogen-limited environments such as the mammalian gut^17^. Specifically, environmental pH modification is a major mechanism shaping the dynamics and metabolism of gut communities cultured *in vitro*^18,19^, such as the production of human health-relevant short chain fatty acids (SCFA)^20,21^. Further, patients with certain GI diseases such as ulcerative colitis can display lower colonic pH than healthy people^22^. This implies that strategies to restore pH homeostasis of the gut could potentially benefit host health^23^.

A reduced butyrate production capability in the human gut microbiome is associated with multiple human diseases^24–26^. Given the substantial inter-individual variability in butyrate production^27^, new strategies to shift the microbiome from low to high butyrate producing states are needed. Predictable modulation of butyrate production via dietary fiber supplementation or the introduction of butyrate-producing species is challenging due to the poorly understood interactions and variation in colonization^28,29^. Previous research has demonstrated that arc operon harboring species *E. lenta* had a substantial effect on the assembly of a 12-member human gut community and positively influenced butyrate production in a 25-member human gut community *in vitro*^30,31^. However, the specific contribution of the *arc* operon to butyrate production and community assembly in human gut communities remains unresolved.

Computational modeling of microbial communities could provide insights into the quantitative role of specialized metabolic pathways on community dynamics and functions^32,33^. While dynamic ecological models such as the generalized Lotka-Volterra (gLV) have been applied to decipher pairwise interactions shaping community assembly^30,31,34,35^, the inferred interaction parameters fail to reveal the modes of interaction. Therefore, computational models that capture key interaction modalities could provide insights into the role of specialized metabolic pathways in the human gut microbiome. For example, a computational model that captures pH-mediated interactions provided insights into the contribution of pH on the assembly of soil microbial communities^36^.

By combining bottom-up design of synthetic human gut communities, computational modeling, and synthetic biology, we investigated the role of the *arc* operon in the human-associated intestinal probiotic strain *Escherichia coli* Nissle 1917 (EcN)^37^. We quantified the effects of the *arc* operon on community assembly, health-relevant metabolite profiles and mammalian gut colonization. We engineered inducible control of the *arc* operon in EcN and investigated the effect of this pathway on the assembly of a defined human gut community both *in vitro* and in the murine gut. By stabilizing the temporal variation in external pH, the *arc* operon reduced variation in community assembly in response to pH perturbations. Further, the presence of the *arc* operon modulated the production of the health-beneficial metabolite butyrate *in vitro* and the abundance of butyrate producers in the mammalian gut. To quantify the role of pH modification of the *arc* operon on community assembly, a dynamic computational model was used to forecast community assembly as a function of monospecies growth and external pH modification. The computational model provided insights into the contribution of pH-mediated interactions shaping community assembly. In sum, we demonstrate that a single specialized metabolic pathway can shape community assembly, species colonization of the mammalian gut and health-relevant metabolite production. Our framework for quantifying the role of specialized metabolic pathways on gut microbiota interactions and functions could be extended to capture other mechanism modalities. In sum, our approach provides a foundation for how to understand and engineer specific molecular mechanisms shaping microbiomes.

## RESULTS

### Arginine dihydrolase pathway increases environmental pH via arginine metabolism

The *arc* operon consists of four genes including arginine deiminase *arcA,* ornithine carbamoyltransferase *arcB,* carbamate kinase *arcC* and ornithine antiporter *arcD* **(Figure 1A)**. The arginine deiminase *arcA* is responsible for the conversion of arginine into citrulline, producing ammonia as a byproduct. The ornithine carbamoyltransferase *arcB* converts citrulline to ornithine and carbamyl phosphate. The carbamate kinase *arcC* further metabolizes carbamyl phosphate to produce more ammonia. Finally, the arginine-ornithine antiporter *arcD*, catalyzes the ATP-independent exchange between arginine and ornithine across the bacterial membrane.

**Figure 1.**
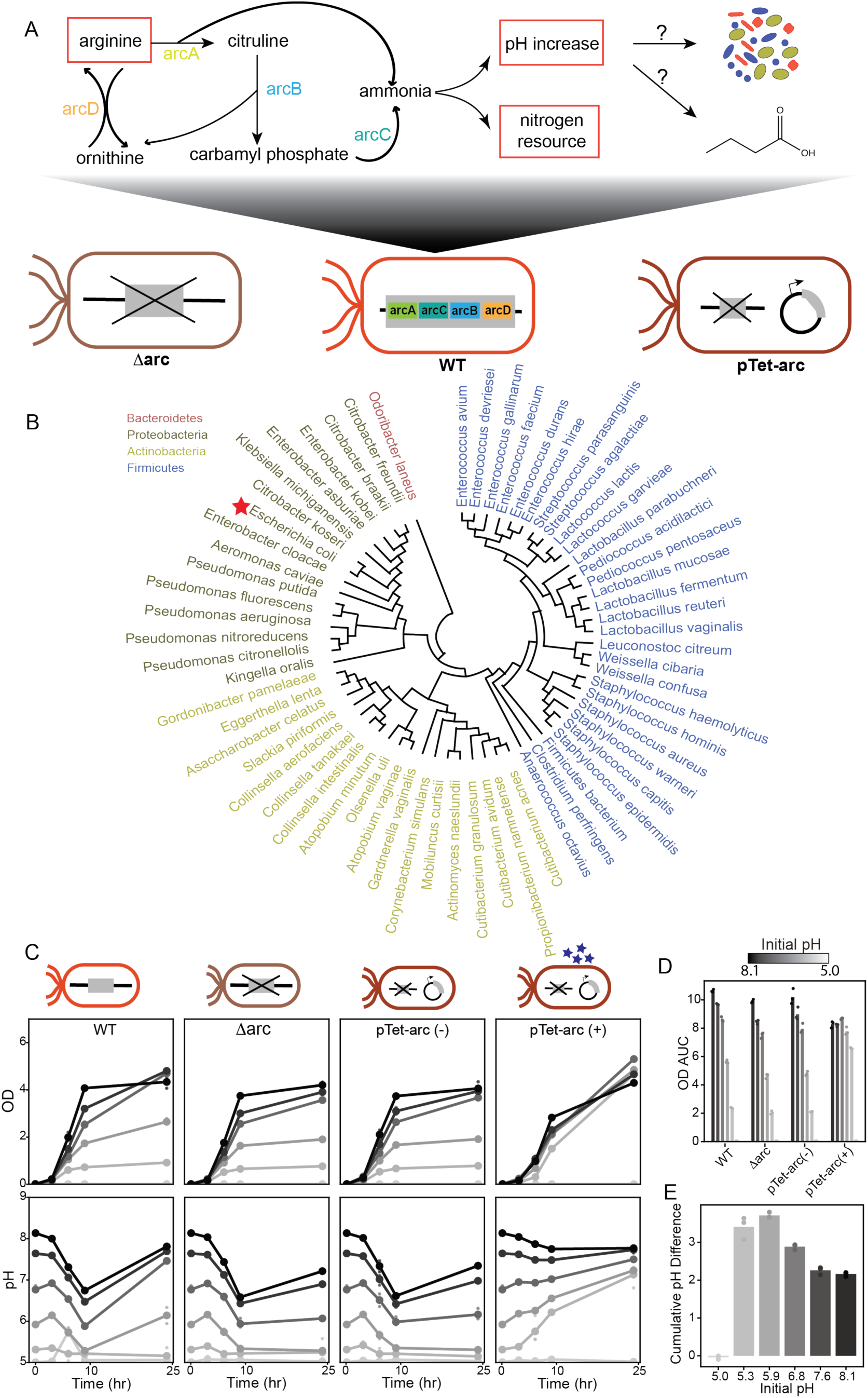
The *arc* operon harbored by a minority of human-associated intestinal bacteria can reduce the variability in *E. coli* Nissle fitness in response to different initial pH environments. **(A)** Schematic of the mechanism of arginine dihydrolase pathway (*arc* operon) and strains used in this study. The *arc* operon in wildtype *E. coli* (EcN) Nissle (WT) encodes four genes for arginine catabolism. The *arc* operon can modulate environmental pH as well as provide a nitrogen source for other constituent community members. EcN strains with *arc* deletion (Δarc) and *arc* under a TetR-inducible promoter on a plasmid (pTet-arc) were constructed for this study. **(B)** Phylogenetic tree of species harboring the arc operon in the human gut microbiome generated by multiple sequence alignment on the 16S region obtained from NCBI taxonomy database^40^. Species color indicates phylum. The species of interest in this study (*E. coli*) is marked by a red star. **(C)** Absolute abundance and supernatant pH of the WT, Δarc, pTet-arc in the absence of aTc (pTet-arc (-)) and pTet-arc in the presence of aTc (pTet-arc (+)) cultured in media with different initial pH ranging from 5.0 – 8.1as a function of time. Smaller sized points indicate 3 biological replicates for each condition. Larger sized points and lines indicate average over all biological replicates. Different initial media pH is marked by different color of line and datapoints, with darker color corresponding to higher initial media pH. **(D)** Total growth quantified by the area under the OD curve (AUC) of absolute abundance measurements for each EcN strain at each initial pH condition. The AUC is calculated by summing the absolute abundance measurements made at each experimental timepoint. The bars represent the average AUC across biological replicates whereas the datapoints represent individual biological replicates (n=3). Colors represent initial pH condition. **(E)** Cumulative pH difference between Δarc and pTet-arc (+) across all timepoints. The measured absolute pH differences between Δarc and pTet-arc (+) at each timepoint are summed to calculate cumulative pH difference for each initial pH condition. The bars represent the average cumulative pH difference across biological replicates whereas the datapoints represent individual biological replicates (n=3).

The prevalence of the *arc* operon has been investigated in oral-associated microbial species and strains of select species^10–12^. To determine the prevalence of the *arc* operon across the human microbiome, we analyzed a metagenome-assembled genome dataset containing 154,723 microbial genomes assembled from 9,428 samples of the human microbiome^38^. The microbiome samples were derived from different body sites in individuals from diverse geographic locations, ages, and lifestyles. Of the 533 total species identified from the annotated database, 63 species (12%) harbor *arcA*, *arcB*, *arC* and *arcD* (**Figure 1B**). The *arc* operon was rarely found in Bacteroidetes, one of the most abundant phyla in human gut microbiome. Additionally, the *arc* operon was harbored by 28 pathogens associated with human gut, oral, or vaginal environments (**Supplementary Data 9).** These results demonstrate that the *arc* operon is a specialized metabolic pathway in the human microbiome due to its low prevalence across species in the dataset.

The probiotic strain *Escherichia coli* Nissle 1917 harbors the *arc* operon, has extensive genetic tools, and has relevance to human health^39^. To evaluate the impact of the *arc* operon on the fitness of EcN, we deleted this operon from the genome of the wildtype EcN (WT) yielding the Δarc strain **(Figure 1A)**. In addition, we constructed pTet-arc by introducing the *arc* operon controlled by an anhydrotetracycline (aTc) inducible promoter onto the genome of Δarc **(Figure 1A)**. Using quantitative reverse transcription polymerase chain reaction (RT-qPCR), the expression of the *arcD* gene **(Supplementary Data 5)** in the *arc* operon of pTet-arc strain was substantially increased in the presence than absence of aTc in EcN monocultures **(Figure S1A)**.

To quantify the effect of the *arc* operon on the growth of EcN, we characterized the growth of each EcN strain in media that varied in initial pH (5.0 to 8.1) and was supplemented with arginine and the presence/absence of aTc (**Figure 1C**). The growth of EcN increased with the initial media pH. The Δarc and pTet-arc strains displayed similar growth and trends in external pH in the absence of aTc (pTet-arc (-)), demonstrating that the pTet promoter was not leaky. The wildtype (WT) and pTet-arc strain induced with aTc (pTet-arc (+)) displayed higher endpoint environmental pH than the Δarc and pTet-arc (-) conditions. Notably, the growth of pTet-arc (+) was similar across a wide range of initial pH values, whereas the other EcN strains exhibited substantially larger growth variability. Specifically, pTet-arc (+) displayed growth in media with an initial pH of 5.3, whereas all other strains displayed low growth (**Figure 1C,D**). In addition, the magnitude of environmental pH differences between the Δarc and pTet-arc(+) strains across all time points displayed a non-monotonic relationship with the initial media pH **(Figure 1E)**. These results highlight that the initial pH influences the pH-modulating activity of the *arc* operon and this activity is maximized at an initial pH of approximately 5.3-5.9.

A previous study demonstrated that the *arc* operon provides an advantage for a pathogenic *E. coli* wildtype strain *in vitro* in the presence of arginine while conferring a colonization disadvantage in the murine gut^13^. To probe the effects of the arc operon more quantitatively on the fitness of EcN in the presence and absence of *arc* operon in EcN, we performed a competition experiment between the WT and Δarc strain. We co-cultured the WT and Δarc strain that each harbor a unique 6-bp barcode on the genome in presence of a range of arginine concentrations and initial pH values. The WT displayed higher abundance than Δarc across all conditions and the magnitude of this difference increased with the initial arginine concentration and media pH **(Figure S1B-D)**. To determine whether this fitness advantage was observed in the mammalian gut, we orally gavaged male germ-free mice (6-8 weeks old) with the co-culture of WT and Δarc strains. The mice received a high arginine diet (5% total arginine, **Supplementary Data 8**) 3 days prior to introduction of the EcN strains. Fecal samples were collected every 2-3 days for quantifying species relative abundance and pH. The relative abundance of the WT was lower than the Δarc strain for most time points for all mice **(Figure S1F-G)**. Therefore, the *arc* operon in EcN confers an advantage *in vitro* and colonization disadvantage *in vivo*, consistent with the trends observed in the pathogenic *E. coli* strain in the previous study^13^. In sum, the expression of the *arc* operon in the pTet-arc strain enabled consistent growth across a wide range of initial pH values and the *arc* operon confers a colonization disadvantage to the WT EcN strain in the mammalian gut.

### The arc operon impacts human gut community assembly and metabolite production in vitro

Since the presence of the *arc* operon is linked to changes in community assembly and butyrate production^31,41^, we investigated the role of the *arc* operon on community assembly and metabolite production in a human gut community. We designed an 8-member human gut community consisting of representatives from the major phyla *Bacteroidetes*, *Firmicutes*, and *Actinobacteria* **(Figure 2A, Supplementary Data 1)** that do not harbor the *arc* operon or other arginine degradation pathways (e.g. Stickland fermentation and the arginase pathway)^42^ to minimize functional redundancy and eliminate arginine competition with EcN. This community contained two prevalent butyrate-producing species (*Anaerostipes caccae* and *Coprococcus comes*), previously shown to contribute substantially to butyrate production in a human gut community^31,43^. In each condition, we introduced an individual EcN strain. The communities were cultured in a chemically defined medium that can support the growth of all species in monoculture^31^. Since expression of the *arc* operon is regulated by arginine availability^44^, the arginine concentration (1%) was chosen to enable activity of *arc* operon and promote community diversity by limiting growth of EcN **(Figure S2)**.

**Figure 2.**
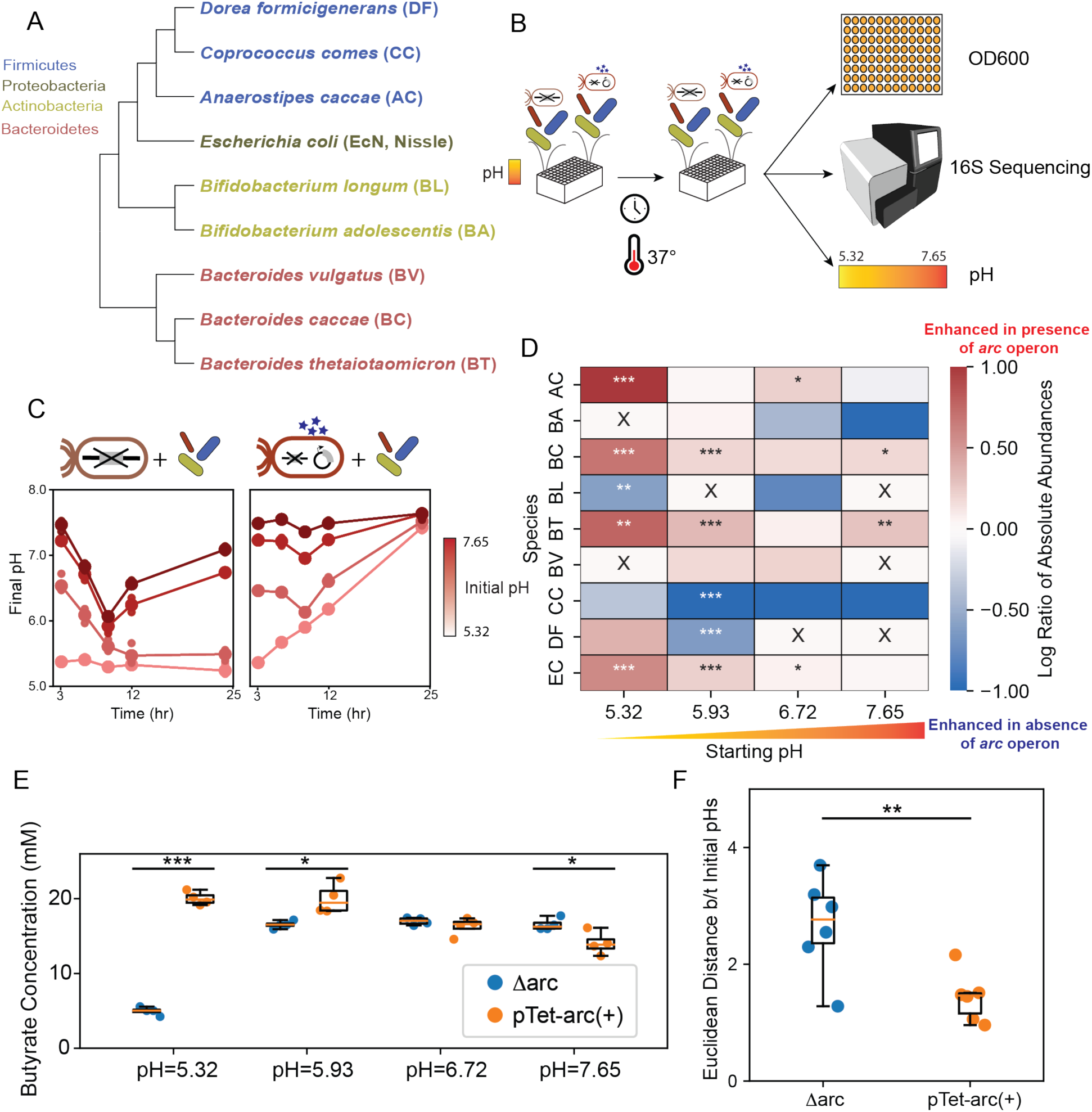
The *arc* operon shapes community assembly and butyrate production and promotes reproducible community assembly in response to pH perturbations. **(A)** Phylogenetic tree of the synthetic human gut microbial community composed of 8 highly prevalent and diverse species generated by multiple sequence alignment on the 16S region obtained from NCBI taxonomy database^40^. *E. coli* Nissle harbors the *arc* operon. **(B)** Schematic of the bottom-up community assembly workflow that includes individual EcN strains at various starting pH. Synthetic communities are cultured in microtiter plates in anaerobic conditions and incubated at 37°C. The absolute abundance of each species is determined by measuring cell density at 600 nm (OD600) and pH. Community composition is determined using multiplexed 16S rRNA sequencing. **(B)** Supernatant pH of community cultures with Δarc (left) or pTet-arc (+) (right) in media with different initial pH values ranging from 5.32 to 7.65. Species were inoculated at an equal initial abundance based on OD600 measurements. Datapoints indicate 4 biological replicates for each experimental condition. Lines indicate average over all biological replicates. Different initial media pH is marked by different color of line and datapoints, with darker color corresponding to higher initial media pH. **(D)** Heatmap of the log of the ratio between species absolute abundance (mean value of biological replicates, n = 4) in pTet-arc (+) and Δarc conditions at various initial media pH at 24 hours. Species relative abundances are determined by multiplexed 16S rRNA sequencing. Red corresponds to higher relative abundance in WT conditions and blue corresponds to higher relative abundance in Δarc conditions. Low growth conditions are marked by an ‘X’. Asterisks represent statistical significance: *P < 0.05, **P < 0.01, ***P < 0.001 according to an unpaired t-test. **(E)** Box plot of butyrate concentration (mM) at 24-hour for each community containing the Δarc or pTet-arc(+) strain at various starting pH. Datapoints indicate 4 biological replicates. Datapoint colors indicate the strain of EcN in the community. An unpaired t-test was performed on each pair of conditions with the same starting pH. Asterisks represent statistical significance: *P < 0.05, **P < 0.01, ***P < 0.001 according to an unpaired t-test. **(F)** Euclidean distance in community structure between all pairs of initial pH conditions at 24-hr for the Δarc (blue) and pTet-arc(+) condition (orange). The Euclidean distances were calculated based on community relative abundance structure between each pair of initial pH condition at 24 hour for a particular strain condition. Data points represent the Euclidean distance calculated for different pairs of initial pH conditions.

Since the pTet-arc (+) and Δarc conditions represent the two extremes of *arc* operon expression **(Figure S1A)**, a comparison of these conditions can provide insights into the impact of the *arc* operon on community assembly and butyrate production **(Figure 2B)**. Notably, high expression of the *arc* operon steered the community towards a very similar endpoint pH despite wide variations in the initial media pH (**Figure 2C**). The *arc* operon also shaped the absolute abundance and relative abundance of different species in the community across different initial pH environments **(Figure 2D, S3)**. Certain shifts in species abundance were consistent with the monoculture growth characterization in response to different initial pH values. For example, species that displayed higher monoculture growth in response to lower initial pH (BL, CC) also were enhanced in the absence of the *arc* operon expression in lower initial pH conditions (5.32, 5.93) at 24 hours **(Figure 2D, S4A)**. Similarly, species that exhibited higher monoculture growth in the presence of higher initial pH (e.g. AC, BC, BT and EcN) also were enhanced in communities with high *arc* operon expression at 24 hours in low pH environments (5.32 and 5.93) **(Figure 2D, S4A)**. Overall, the growth profiles of each species in the presence and absence of the community displayed similar trends except for DF, which grew poorly in monoculture **(Figure S4A)**. Finally, except initial pH condition 5.32, the impact of the *arc* operon on the growth of the butyrate producer AC varied as a function of time. In the initial pH conditions 5.93, 6.72 and 7.65, AC was enhanced at earlier times and was then inhibited or displayed no significant change at later times **(Figure S4C-E)**. The time-dependent changes in the impact of *arc* operon on the growth of AC potentially arise due to a diauxic shift in metabolism of AC or inter-species interactions. This demonstrates that the presence of the *arc* operon can shape transient and sustained shifts in species abundance as a function of time.

High expression of the *arc* operon (pTet-arc(+)) impacted the production of butyrate by the synthetic human gut community **(Figure 2E).** The physiological pH range of the human caecum and large intestine which contains the largest number of gut bacteria is 5-7^45,46^. Within this range, butyrate was significantly higher in the presence of pTet-arc (+) than in the absence of the *arc* operon (Δarc). This trend was consistent with a growth enhancement of AC (**Figure S5A**). By contrast, the absolute abundance of CC was lower in pTet-arc (+) conditions at initial pH of 5.93 **(Figure S5B)**, highlighting the differential effects of the *arc* operon on diverse butyrate producing bacteria. At the highest initial pH (pH=7.65), the reverse trend was observed where the absence of the *arc* operon yielded significantly but moderately higher butyrate. The total butyrate producer abundance displayed a moderate positive correlation with the endpoint butyrate concentration **(Figure S5C).** The lactate and butyrate concentrations were inversely related at an initial pH=5.32 **(Figure 2E**, **S6)**. In sum, this implies that the *arc* operon enhanced the abundance of AC and its ability to convert lactate to butyrate in this condition **(Figure S5A)**. While AC can convert acetate into butyrate^47^, acetate can also be released by other species in the community masking its utilization **(Figure S6A)**^48^. These results highlight the differential effects of the *arc* operon on health-relevant metabolite profiles of human gut communities.

To assess the effect of the *arc* operon on the sensitivity of community assembly to initial pH perturbations, we evaluated the pairwise Euclidean distances of community composition at the endpoint across different initial pH conditions. The variance in the Euclidean distance distribution was substantially lower in communities with high *arc* operon expression than in the absence of the *arc* operon (**Figure 2F**). This implies that the *arc* operon can stabilize the assembly of the community in response to external pH perturbations (**Figure 2C**). In sum, the *arc* operon in EcN can promote reproducible community assembly in response to variations in initial pH and shift the metabolic states of human gut communities from low to high butyrate production in physiologically relevant pH ranges.

### A dynamic computational model reveals the contributions of external pH to community assembly

To quantitatively understand the pH-dependent effects of the *arc* operon on *in vitro* community assembly, the following system of ordinary differential equations was used to model species and pH dynamics^36^,

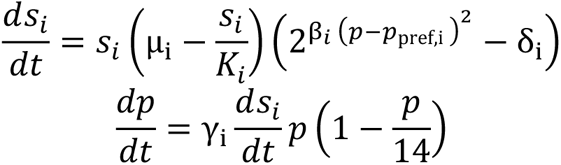

where *S_i_* is the abundance (OD600) of species *i*, and *p* represents pH. Model parameters include growth rate µ_i_, carrying capacity *K_i_*, preferred pH, p_pref,i_, sensitivity to deviation from the preferred pH, β_i_, and constant γ_i_ that relates a change in species growth to a change in pH. The model captures growth rate maximization at a preferred pH. Consistent with our experimental data, the model displays a positive growth rate when pH is close to preferred and becomes negative when pH deviates substantially from preferred **(Figure S7, S8).** The model assumes that the contribution of each species to environmental pH is linearly related to growth rate, consistent with the similar temporal trends in pH and growth **(Figure S7, S8)**. We fit the model to monospecies time-series measurements of OD600 and pH over a 24-hour period. Species were cultured in the presence of six different initial pH values ranging from 5 to 8.

We used an approximate Bayesian parameter estimation approach referred to as variational inference to optimize a Gaussian approximation of the parameter posterior distribution and the precision of a zero-mean Gaussian parameter prior and zero-mean Gaussian measurement noise **(see Supplementary Text)**. To validate the predictive performance of the model, we performed leave-one-out cross-validation, where data from all except one initial pH condition were used for training and the remaining pH condition was held-out for testing. The process was repeated until all initial pH conditions were subject to held-out testing. Model prediction performance was evaluated using the Pearson correlation coefficient between predicted and measured species OD600. The model accurately predicted the growth of most species with the exception of BC, BT, BV and DF, which displayed no or moderate growth in only a subset of conditions **(Figure S4A, S8, S9)**. The model displayed a moderate fit to the non-monotonic temporal changes in pH of WT and pTet-arc (+) **(Figure S8 C,E)**. This is consistent with the model’s inability to capture the diauxic growth of EcN strains with *arc* operon, which could yield a time-dependent change of the effect of EcN on pH **(Figure S8C-E)**^49^.

Monoculture fitting of the pH model indicates that certain species prefer lower pH (BL), whereas others prefer higher pH (AC) **(Figure 3A, S4A)**. The effects of species on environmental pH also varied (e.g. BA, BL, BT, BV, and CC decrease environmental pH whereas BC increases environmental pH) **(Figure 3A)**. The model fit to monoculture data was used to predict species growth dynamics in the community. The model prediction of community assembly displayed similarities to experimental data in the presence of lower initial pH **(Figure S10A).** The prediction of species presence or absence across different initial pH conditions was largely consistent with most experimental conditions except for the highest initial pH condition **(Figure S10B).** In addition, the model consistently overpredicted AC **(Figure S10A).**

**Figure 3.**
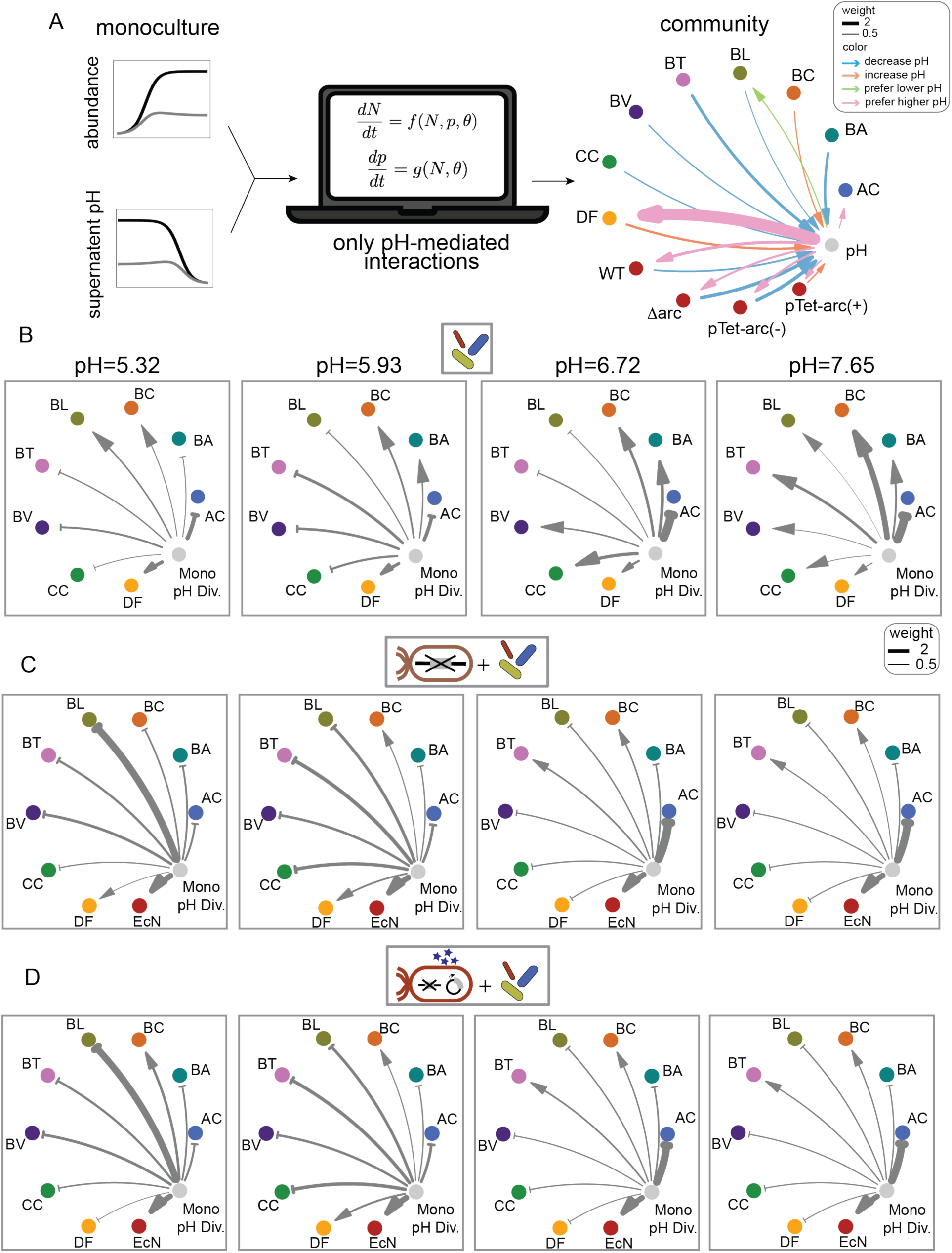
Investigating the contribution of pH-mediated gut microbiota interactions on community assembly in different pH environments. **(A)** Computational modeling framework for predicting community assembly. The pH model is a coupled set of ordinary differential equations that captures species growth and pH modification. Model parameters were estimated by fitting to monoculture data. The inferred interaction network of species and pH (right) highlights the contributions of species to environmental pH and the environmental pH of maximum growth for each species. Edges from pH to species represent species maximized growth at higher (pink) or lower (green) environmental pH, calculated by *p_pref,i_* − 6.5, where *p_pref,i_* represents the preferred pH parameter for species *i* from the model. Edges from species to pH represent effect of species growth to increase (orange) or decrease (blue) environmental pH, calculated by 20 ∗ *γ_i_*, where *γ_i_* represents effect of species *i* growth on pH and a factor of 20 was introduced for visualization. **(B), (C), (E)** monoculture-pH-divergence networks of cocultures of gut community (B), community with Δarc (C), and community with pTet-arc (+) (E) at different initial pH. Model was used to predict the assembly for each coculture across different initial pH conditions. The absolute values of monoculture-pH-divergence are calculated by the root mean squared error between model prediction and experimental data for each species in each coculture, averaged over all timepoints. The signs of monoculture pH divergence are determined by the sign of the difference between model prediction and experimental data. Species that are over or under predicted by the pH model are represented by blunted or arrows, respectively. Arrow width represent the strength of monoculture pH divergence, multiplied by a factor of 10 for visualization. Arrow width values greater than 3 are rounded to 3 for visualization.

To provide deeper insights into changes in pH mediated interactions in monoculture versus community contexts, we define a species-specific metric for interactions that deviate from the monoculture informed model (i.e. “monoculture pH divergence”). Monoculture pH divergence is defined as the root mean squared error (RMSE) between predicted and measured species relative abundance across all timepoints in each initial pH condition (**Figure 3B, C, D**). Larger monoculture pH divergence in communities suggest interactions not captured by monoculture pH dynamics, which could arise from either non-pH-mediated interactions (e.g. resource competition, cross-feeding or anti-microbial effects) or pH-mediated interactions that deviate from monoculture behaviors (e.g. changes in metabolic niche in the community than monoculture which alters the impact of a given species on pH). Species underpredicted by the model are influenced by growth promoting interactions that deviate from monoculture pH effects and species that are overpredicted by the model are influenced by growth inhibitory interactions that deviate from monoculture effects **(Figure 3B)**. For example, in monoculture, DF failed to grow in all conditions **(Figure S8)** but displayed growth in most conditions in the community, highlighting the role of growth promoting inter-species interactions not captured by the model. By contrast, AC is consistently overpredicted in the community **(Figure 3B),** suggesting missing inhibitory interactions with AC that are not captured by the model.

To provide insights into the extent of interactions captured by monoculture pH dynamics in the presence of high expression of the *arc* operon (pTet-arc(+)) or absence of the *arc* operon (Δ arc), we evaluated the difference between the pH model prediction and experimental measurements of the human gut community containing individual EcN strains **(Figure S11)**. The pattern of species presence/absence in the model and experimental data was more consistent in the presence of Δarc than pTet-arc(+) **(Figure S12A,B)**. The model accurately predicts the non-monotonic trends in the Shannon diversity index as a function of initial environmental pH **(Figure S12C)**. However, the pH dynamics were not well-captured by the model at later time points **(Figure S11)**. This deviation between model and experiment may arise from the poor model fit to the pH dynamics in the EcN monocultures, which impacted community assembly as the highest abundance species.

The network visualization of monoculture pH divergence can reveal how interactions beyond monoculture pH vary across different initial pH environments and communities **(Figure 3B, C, D)**. Overprediction and underprediction of species by the pH model suggests the presence of growth inhibitory and growth promoting inter-species interactions that deviate from monoculture pH effects, respectively. Substantial changes in monoculture pH divergence for a given species across conditions suggests that the species interaction modalities vary across environments (i.e. initial pH or community context). Certain species (e.g. DF, BC, BA) have low monoculture pH divergence across initial pH conditions in the presence of EcN **(Figure 3C, D)**. This implies that pH mediated interactions can explain a large fraction of the variance in their growth dynamics in the community. Certain species displayed altered monoculture pH divergence in the presence of EcN versus absence of EcN (e.g. BL in the initial pH 5.32 condition and BA in the initial pH 5.93 condition) **(Figure 3B, C, D),** suggesting that their interaction modalities were altered in the presence of EcN. By contrast, most species displayed similar monoculture pH divergence in the presence and absence of the *arc* operon **(Figure 3C, D)**, despite changes in species growth dynamics in the community between these conditions **(Figure S11)**. Overall, our results suggest that the *arc* operon did not substantially alter the interaction modalities of most species in the community.

To provide insights into the contribution of pH modification on community assembly across conditions, we evaluated whether changes in pH due to *arc* operon activity displayed an informative relationship with changes in community composition. To this end, we evaluated the mapping between the Euclidean distance in community composition and the difference in pH in the presence (pTet-arc(+)) and absence (Δarc) of the *arc* operon at each time point. A positive correlation implies that alterations in community composition display an informative relationship with the pH modification activity of the *arc* operon. A positive correlation was observed for the initial pH=5.93 condition (Spearman *ρ* = 0.72 with p-value =0.0004) **(Figure S12D),** suggesting that *arc* operon activity shaped community assembly via pH modification in this environment. In sum, our monoculture-informed pH model provided insights into the quantitative contributions of monoculture pH dynamics on community assembly.

### Investigating the effects of the arc operon on community assembly and SCFA production in the murine gut

We investigated the effects of the *arc* operon on community assembly and health-relevant metabolite profiles in the mammalian gut by colonizing male gnotobiotic mice (6-8 weeks old) with a human gut community that harbored or did not harbor the *arc* operon. Mice were fed a high arginine diet **(Supplementary Data 8)** three days prior to oral gavage with the community cultures. This diet contained 5% arginine, which is higher than an arginine concentration previously shown to elevate the arginine in the murine colon which yielded an altered microbiome composition^50,51^. Mice were orally gavaged with a culture containing equal species proportions of the 8-member community. After one week, groups of mice received the EcN WT, Δarc or no EcN strain via oral gavage. To characterize the changes in community composition over time, fecal samples were collected every 2-3 days for measurement of pH^52,53^ and 16S rRNA gene sequencing. After two weeks, mice were sacrificed and cecal contents were collected for measurement of health-relevant metabolites, pH^52,53^ and community composition based on 16S rRNA gene sequencing **(Figure 4A)**. Using bacterial RNA extracted from cecal contents, the WT *arc* operon displayed higher expression than the pTet-arc(-) *in vitro* control, suggesting that the *arc* operon was expressed in the caecum **(Figure S1A)**.

**Figure 4.**
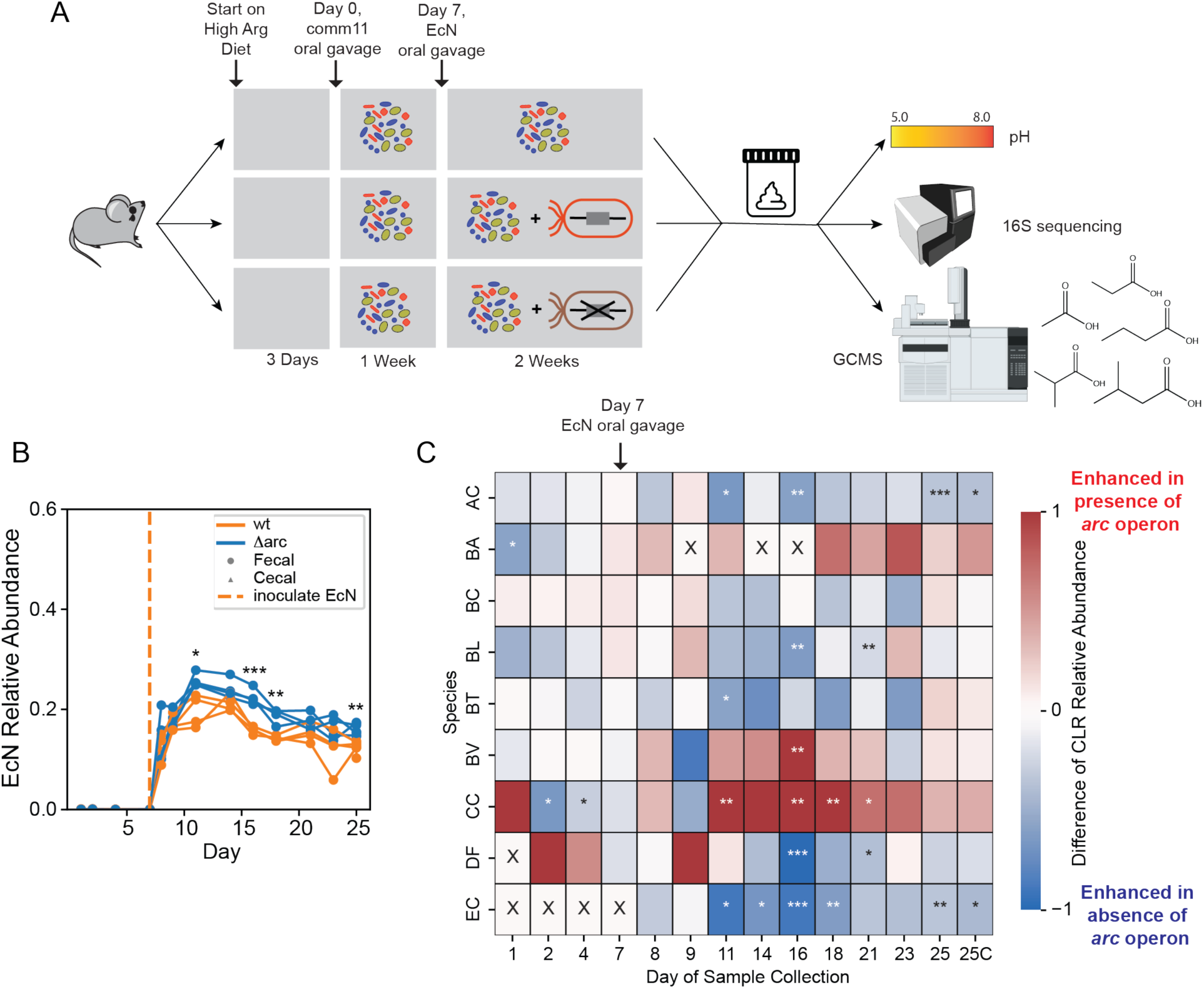
The *arc* operon impacts community assembly and species colonization of the mammalian gut. **(A)** Schematic of communities characterized in germ-free mice. Mice were tagged and placed on customized high arginine diet 3 days before oral gavage of human gut communities. Fecal samples were collected every two to three days and characterized via multiplexed 16S rRNA sequencing and fecal sample pH. After 1 week of colonization, WT or Δarc strains of EcN were introduced by oral gavage. After 2 weeks, mice were sacrificed and the community composition, pH and metabolite concentrations were measured using the cecal contents. **(B)** EcN relative abundance in fecal and cecal samples in WT and Δarc conditions. EcN relative abundances are determined by multiplexed 16S rRNA sequencing on fecal samples and cecal samples. Datapoints indicate biological replicates. Lines connect each of 4 biological replicates collected from different mice. EcN inoculation is marked by orange dashed line on Day 7. Asterisks represent statistical significance: *P < 0.05, **P < 0.01, ***P < 0.001 according to an unpaired t-test. **(C)** Heatmap of the difference in species relative abundance (mean value of biological replicates, n = 3) in the presence of WT or Δarc at various timepoints. Red corresponds to higher relative abundance in the presence of the WT strain and blue corresponds to higher relative abundance in the presence of the Δarc strain. EcN inoculation is marked on Day 7. Asterisks represent statistical significance: *P < 0.05, **P < 0.01, ***P < 0.001 according to an unpaired t-test.

The relative abundance of the WT and Δarc EcN strains increased at the same rate and saturated at day 4 **(Figure 4B)**. The Δarc strain colonized mice at a moderately but significantly higher abundance than the WT strain over several timepoints, consistent with its higher abundance in competition with the WT strain in an *E. coli* monocolonized mouse **(Figure S1)**. Further, both EcN strains converged to approximately 20% in the presence of the human gut community **(Figure 4B, S13A-B)**. Certain species were enriched in the presence of the *arc* operon (WT condition) *in vivo* (CC, BC), whereas the colonization of other species was reduced (AC, DF) at different timepoints **(Figure 4C)**. The enhancement or inhibition of certain species in the presence versus absence of the *arc* operon were consistent between *in vitro* (initial pH=5.32) and *in vivo* (e.g. BL) whereas other species displayed inconsistent trends (e.g. AC, EcN) **(Figure 2D)**. The effects of the *arc* operon on BC and DF were transient, whereas the changes in abundance of the butyrate producers CC and AC were sustained over multiple timepoints **(Figure S4B-E)**. These results highlight the key role of the mammalian gut in shaping community dynamics that display both consistent and inconsistent trends with *in vitro* data.

The fecal and cecal pH were not significantly different between groups **(Figure S13D)**, indicating that the *arc* operon did not have a substantial effect on the pH of the fecal and cecal samples. Further, measurements of the health-relevant metabolites butyrate, propionate, isobutyrate, acetate and isovalerate in cecal contents were not significantly different in the presence (WT) and absence ( Δ arc) of the *arc* operon at the endpoint **(Figure S14).** The abundance of most species (AC, BL, BV, CC, DF, EcN) in the community were different between conditions on Day 16, whereas only AC and EcN were altered between conditions at the time of cecal content harvest. Consistent with this trend, the difference in relative abundance of Δarc versus the WT was largest at Day 16. Therefore, it is possible that health-relevant metabolite production may exhibit differences in the presence/absence of the *arc* operon.

## DISCUSSION

We explored the controllability of human gut community assembly and health-relevant metabolite production by manipulating the expression of a specialized metabolic pathway in a probiotic bacterial species *E. coli* Nissle. By integrating synthetic biology and bottom-up community construction, we showed that the expression of the *arc* operon can promote robustness of communities to pH perturbations and increase butyrate production at physiologically relevant pH ranges *in vitro*. Using a computational model that captures monoculture-informed pH mediated interactions, we uncovered the extent of other interaction modalities not captured by the model on community assembly and showed that the *arc* operon does not alter the interaction modalities for most species during *in vitro* community assembly **(Figure 3C,D)**.

Community robustness is defined as the ability to maintain composition and/or target functions in response to environmental perturbations^54^. Enhancing the robustness of the human gut microbiome to perturbations holds therapeutic potential. For instance, the pH in caecum and colon has been shown to be lower in active patients with ulcerative colitis (4.7) than inactive patients (4.9–5.5) and healthy volunteers (7.2–7.5)^22^. Therefore, promoting community robustness to pH perturbations could reduce the variation in gut microbiome composition and function in certain disease states. The *arc* operon expression modulates environmental pH and community dynamics more at an initial lower pH (pH=5.32) than at higher initial pH (pH=7.32) **(Figure 2C, D)**. This implies that the *arc* operon could potentially promote robustness of gut microbiota to pH perturbations to a larger degree in ulcerative colitis where environmental pH is generally lower than in a healthy state. Therefore, an avenue of future research is exploring the effects of the *arc* operon on community dynamics and health-relevant metabolites in the presence of inflammation and dysbiosis in a murine model of colitis.

A depletion of butyrate producing bacteria in the human gut microbiome has been associated with a wide range of human diseases^55,56^. Developing new strategies to enhance butyrate production are needed due to the large inter-individual variation observed in butyrate production in response to dietary fiber interventions^57^. While the *arc* operon enhanced butyrate production at physiologically relevant pH values, the *arc* operon displayed differential effects on diverse butyrate producing bacteria. In addition, butyrate production was not altered in the murine gut at the measured time point in the presence versus absence of the *arc* operon. The disparate effects of the *arc* operon on butyrate production *in vitro* and *in vivo* could be attributed to the following possibilities: (1) the host may dominate pH homeostasis, yielding negligible effects of the *arc* operon, (2) changes in butyrate production may be transient mirroring the transient changes in the abundance of CC **(Figure 4C)** and were therefore not observed, (3) the *arc* operon has variable or insufficiently low expression in WT EcN *in vivo* to yield environmental impact or (4) changes in the rate of uptake of butyrate by the host could mask potential differences in the presence/absence of the *arc* operon. Synthetic promoters controlling the *arc* operon would allow for more consistent and tunable expression in the mammalian gut. Antibiotic-free selection mechanisms for stable plasmid maintenance may be useful for achieving high expression of the *arc* operon on a multi-copy plasmid^58^.

Widely used ecological models such as the generalized Lotka-Volterra fail to capture specific interaction modalities^31,41^. The computational workflow assuming only pH-mediated interactions trained on monoculture kinetics can reveal the contribution of this mechanism community assembly. Previous studies in the same chemically defined media have identified strong resource competition with AC for the limiting pool of sugars^31^, which could contribute to the consistently negative interactions shaping AC that could not be explained by the monoculture informed pH model **(Figure 3B, D, E)**. Monoculture pH divergence did not change substantially in the presence/absence of *arc* operon expression despite changes in community dynamics **(Figure 3D, E)**. This implies that *arc* operon expression did not change the interaction modalities for most species in community assembly. The deviation between model prediction and experimental data observed in all communities suggest the presence of substantial interactions beyond pH informed by monoculture on community assembly. These mechanisms could include resource competition, metabolite cross-feeding, and antimicrobial production. A similar approach could be employed to guide the mechanistic investigation of other types of interaction modalities including effects of toxin production^59^, by constructing a model that captures this mechanism^60^. A deeper understanding of the driving molecular mechanisms could reveal new mechanistic control parameters for manipulating the microbiome to our benefit.

The *arc* operon may influence community dynamics and functions via release of nitrogen. Nitrogen limited environments, like the mammalian gut^61^, can limit microbial growth^62^. In the mammalian gut, the *arc* operon may have a larger impact on community assembly via nitrogen release as opposed to external pH modulation. To investigate the contribution of nitrogen release via *arc* operon activity, nitrogen availability could be measured in the presence and absence of *arc* operon activity using the carbon-to-nitrogen (C/N) ratios of gut microbes and those of digestible material (derived from digesta or faeces)^63^. An elevated C/N ratio in resources (based on the digestible material) relative to the C/N ratio of consumers (based on the microbes) would suggest large amounts of food are being processed to gain sufficient nitrogen for microbial growth, suggesting nitrogen limitation. Release of nitrogen via the *arc* operon could increase the C/N ratios of microbes to digestible material. Characterizing the C/N ratio of microbes and digestible material in the presence and absence of the *arc* operon could provide key insights into the observed differences between the effect of the *arc* operon *in vitro* and *in vivo*. For example, the *arc* operon had differential effects on the abundance of EcN, AC and CC *in vitro* versus *in vivo* **(Figure 2D, 4B-C, S1F**).

A major goal for microbiome engineering is to discover and exploit molecular control knobs to steer microbiomes to desired states. Specialized metabolic pathways, which can provide key resources for specific gut bacteria or shape global environmental parameters, could be exploited to shape functions of gut microbiota. By integrating mechanistic computational modeling, synthetic biology and bottom-up community assembly, we revealed the extent to which a single mechanism of interaction contributed to community assembly. While we focused on a single metabolic pathway, high-throughput genetic manipulation approaches could also be used to discover and rank the influence of diverse molecular pathways on target community functions^64–66^. Our strategy to quantitatively determine the contribution of metabolic pathways could be applied to the pathway hits to identify novel control knobs for steering microbiomes to desired states.

## METHOD

### Strain maintenance and culturing

All aerobic culturing was carried out in 37°C incubator with shaking. All anaerobic culturing was carried out in an anaerobic chamber with an atmosphere of 2 ± 0.5% H2, 15 ± 1% CO2 and balance N2. All prepared media and materials for anaerobic experiments were placed in the chamber at least overnight before use to equilibrate with the chamber atmosphere. The permanent stocks of each strain used in this paper were stored in 25% glycerol at −80 °C. Batches of single-use glycerol stocks were produced for each strain by first growing a culture from the permanent stock in ABB media **(Supplementary Data 2)** to stationary phase, mixing the culture in an equal volume of 50% glycerol, and aliquoting 400 μL into Matrix Tubes (ThermoFisher) for storage at −80 °C. Quality control for each batch of single-use glycerol stocks included Illumina sequencing of 16S rDNA isolated from pellets of the aliquoted mixture to verify the identity of the organism. For each experiment, precultures of each species were prepared by thawing a single-use glycerol stock (SUGS) and adding 100 μL of SUGS to 5 mL fresh YBHI media **(Supplementary Data 2)** for incubation at 37 °C for 16 hours **(Supplementary Data 1)**. ^31^

### Plasmid and strain construction

All engineered strains and plasmids can be available upon request to corresponding author. PCR amplifications were performed using Phusion High-Fidelity DNA polymerase (New England Biolabs) and oligonucleotides for cloning were obtained from Integrated DNA Technologies. Standard cloning methods were used to construct plasmids. Small scale electroporation was used for plasmid transformation into *E. coli*. The background strain was grown to an OD of 0.3 and then placed on ice for 10 minutes. 1mL of the cell culture was then washed with 10% glycerol two times, with the cells spun down, supernatant drawn out, and resuspended each time. After final wash, cells were resuspended in 100μL 10% glycerol and 1μg PCR product and transferred to 1mm gap electro-cuvette. Cells were electroporated using the the EC1 protocol of BioRad Micropulser Electroporator, then recovered in LB plus appropriate antibiotic for 1 hour. Transformants were selected on LB agar plates with appropriate antibiotic resistance and single colonies were stored.

The native *arc* operon (consisting of genes *arcA, arcC, arcB, arcD,* and *arcR*) was deleted from the genome of WT by replacement with a Kananycin FRT construct amplified from pKD4^67^ **(Supplementary Data 4)** using primers oYL36 and oYL37 **(Supplementary Data 5)**, with Lambda red expression provided by pMP11^68^ **(Supplementary Data 4)**. Transformants were selected on antibiotic selective plates and verified by colony PCR. The kanamycin cassette was then removed from the genome by transforming a FLP recombinase vector pFLP2^69^. Successful Kanamycin removal was screen for the absence of Kanamycin resistance and verified by colony PCR. Finally, the pFLP2 plasmid was cured by growth in 5% sucrose. Final colonies were screened for absence of Carbenicillin resistance. The resulting strains is Δarc.

For inducible expression of the *arc* operon, we replaced the RFP region of pBbA2a-RFP plasmid^70^ **(Supplemental Data 4)** with the native *arc* operon without regulatory region (consisting of genes arcA, arcC, arcB, and arcD) from WT *E. coli* strain. The *arc* operon was amplified by oYL19 and oYL20, and the pBbA2a-RFP backbone was amplified by oYL17 and oYL18 **(Supplemental Data 5)**. Standard PCR and Gibson assembly was performed to yield pTet-arcACBD **(Supplemental Data 4)**. Finally, strain pTet-arc **(Supplemental Data 1)** was cloned by transforming plasmid pTet-arcACBD into strain Δarc.

### Monoculture and community culturing dynamic growth quantification

Each species’ preculture was diluted to an OD600 of 0.01 (Tecan F200 Plate Reader) in a defined media named DM38^31^ designed in a previous study to enable growth of all species in this study. Precultures were then aliquoted into three or four replicates of 1.5 mL each in a 96 Deep Well (96DW) plate and covered with a semi-permeable membrane (Diversified Biotech) and incubated at 37 °C without shaking. At each time point, samples were mixed by pipetting up and down 5 times before transferring 200uL to 96 well microplate for measurement of OD and supernatant pH were measured by the plate reader (Tecan F200 Plate Reader). Cell pellet was saved at each timepoint for species abundance quantification at −80°C. For pH measurement, phenol red solution was diluted to 0.005% weight per volume in MilliQ water. Bacterial supernatant (20 μL) was added to 180 μL of phenol red solution, and absorbance was measured at 560 nm using the plate reader. A standard curve was produced by fitting the Henderson–Hasselbach equation to fresh media with pH standards ranging between 3 and 11 measured using a standard electro-chemical pH probe (Mettler-Toledo). We used the following equation to map the pH values to the absorbance measurements.

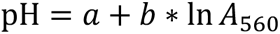

The parameters a and b were determined using a linear regression between pH and the log term for the standards in the linear range of absorbance with A representing the absorbance of each condition.

### Genomic DNA extraction from cell pellets

Genomic DNA was extracted from cell pellets using a modified version of the Qiagen DNeasy Blood and Tissue Kit protocol. First, pellets in 96DW plates were removed from −80 °C and thawed in a room temperature water bath. Each pellet was resuspended in 180 μL of enzymatic lysis buffer (20 mM Tris–HCl (Invitrogen), 2 mM sodium EDTA (Sigma-Aldrich), 1.2% Triton X-100 (Sigma-Aldrich), 20 mg/mL lysozyme from chicken egg white (Sigma-Aldrich)). Plates were then covered with a foil seal and incubated at 37 °C for 30 min with orbital shaking at 600 RPM. Then, 25 μL of 20 mg mL−1 Proteinase K (VWR) and 200 μL of Buffer AL (QIAGEN) were added to each sample before mixing with a pipette. Plates were then covered by a foil seal and incubated at 56 °C for 30 min with orbital shaking at 600 RPM. Next, 200 μL of 100% ethanol (Koptec) was added to each sample before mixing and samples were transferred to a nucleic acid binding (NAB) plate (Pall) on a vacuum manifold with a 96DW collection plate. Each well in the NAB plate was then washed once with 500 μL buffer AW1 (QIAGEN) and once with 500 μL of buffer AW2 (QIAGEN). Samples were then eluted into a clean 96DW plate from each well using 110 μL of buffer AE (QIAGEN) preheated to 56 °C. Genomic DNA samples were stored at −20 °C until further processing.

### Sequencing library preparation

Genomic DNA concentrations were measured using a SYBR Green fluorescence assay and then normalized to a concentration of 1 ng μL−1 by diluting in molecular grade water using a Tecan Evo Liquid Handling Robot. First, genomic DNA samples were removed from −20 °C and thawed in a room temperature water bath. Then, 1 μL of each sample was combined with 95 μL of SYBR Green (Invitrogen) diluted by a factor of 100 in TE buffer (Integrated DNA Technologies) in a black 384-well microplate. This process was repeated with two replicates of each DNA standard with concentrations of 0, 0.5, 1, 2, 4, and 6 ng μL−1. Each sample was then measured for fluorescence with an excitation/emission of 485/535 nm using a Tecan Spark plate reader. Concentrations of each sample were calculated using the standard curve and a custom Python script was used to compute the dilution factors and write a worklist for the Tecan Evo Liquid Handling Robot to normalize each sample to 1 ng μL−1 in molecular grade water. Samples with DNA concentration <1 ng μL−1 were not diluted. Diluted genomic DNA samples were stored at −20 °C until further processing.

Amplicon libraries were generated from diluted genomic DNA samples by PCR amplification of the V3–V4 of the 16S rRNA gene using custom dual-indexed primers (Supplementary Data 3) for multiplexed next-generation amplicon sequencing on Illumina platforms (Method adapted from Venturelli et al. Mol. Syst. Biol., 2018). Primers were arrayed in skirted 96-well PCR plates (VWR) using an acoustic liquid handling robot (Labcyte Echo 550) such that each well received a different combination of one forward and one reverse primer (0.1 μL of each). After liquid evaporated, dry primers were stored at −20 °C. Primers were resuspended in 15 μL PCR master mix (0.2 μL Phusion High Fidelity DNA Polymerase (Thermo Scientific), 0.4 μL 10 mM dNTP solution (New England Biolabs), 4 μL 5× phusion HF buffer (Thermo Scientific), 4 μL 5 M Betaine (Sigma-Aldrich), 6.4 μL Water) and 5 μL of normalized genomic DNA to give a final concentration of 0.05 μM of each primer. Primer plates were sealed with Microplate B seals (Bio-Rad) and PCR was performed using a Bio-Rad C1000 Thermal Cycler with the following program: initial denaturation at 98 °C (30 s); 25 cycles of denaturation at 98 °C (10 s), annealing at 60 °C (30 s), extension at 72 °C (60 s); and final extension at 72 °C (10 min). 2 μL of PCR products from each well were pooled and purified using the DNA Clean & Concentrator (Zymo) and eluted in water. The resulting libraries were sequenced on an Illumina MiSeq using a MiSeq Reagent Kit v3 (600-cycle) to generate 2 × 300 paired-end reads.

### Bioinformatic analysis for quantification of species abundance

Sequencing data were demultiplexed using Basespace Sequencing Hub’s FastQ Generation program. Custom python scripts were used for further data processing (method adapted from Venturelli et al. Mol. Syst. Biol., 2018)16. Paired end reads were merged using PEAR (v0.9.10)83 after which reads without forward and reverse annealing regions were filtered out. A reference database of the V3–V5 16S rRNA gene sequences was created using consensus sequences from next-generation sequencing data or Sanger sequencing data of monospecies cultures. Sequences were mapped to the reference database using the mothur (v1.40.5)84 command classify.seqs (Wang method with a bootstrap cutoff value of 60). Relative abundance for each species within a sample was calculated as the read count mapped to each species divided by the total number of reads for each sample. Absolute abundance of each species was calculated by multiplying the relative abundance by the OD600 measurement for each sample. Samples were excluded from further analysis if >1% of the reads were assigned to a species not expected to be in the community (indicating contamination) or if they had <1000 total reads and OD600 > 0.1 (indicating that there were insufficient reads for analysis and this was not due to lack of community growth).

### HPLC quantification of organic acids from *in vitro* samples

Supernatant samples (200uL) were thawed in a room temperature water bath before addition of 2 μL of H_2_SO_4_ to precipitate any components that might be incompatible with the running buffer. The samples were then centrifuged at 3500 × rpm for 10 min and then 150 μL of each sample was filtered through a 0.2 μm filter using a vacuum manifold before transferring 70 μL of each sample to an HPLC vial. HPLC analysis was performed using a Shimadzu HPLC system equipped with a SPD-20AV UV detector (210 nm). Compounds were separated on a 250 × 4.6 mm Rezex© ROA-Organic acid LC column (Phenomenex Torrance, CA) run with a flow rate of 0.2 mL min^−1^ and at a column temperature of 50 °C. The samples were held at 4 °C prior to injection. Separation was isocratic with a mobile phase of HPLC grade water acidified with 0.015 N H_2_SO_4_ (415 µL L−1). Two standard sets were run along with each sample set, one before all samples and one after all samples. Standards were 100, 20, and 4 mM concentrations of butyrate, succinate, lactate, and acetate, respectively. The injection volume for both sample and standard were 25 µL. The resultant data was analyzed using Shimadzu LabSolutions software package with peaks manually reintegrated if necessary.

### Gnotobiotic mouse experiments

All germ-free mouse experiments were performed following protocols approved by the University of Wisconsin-Madison Animal Care and Use Committee. Two diets were used in this experiment: regular diet with lower arginine concentration (Chow diet, Purina, LabDiet 5021) and high arginine diet (Envigo, TD.210715). All strains were grown at 37 °C anaerobically in YBHI media (Acumedia, Bacto, and Sigma-Aldrich) for 16hrs. All strains for oral gavage were mixed in equal proportions based on OD600 and transferred on ice prior to oral gavage. We used 8-week old C57BL/6 gnotobiotic male mice (wild-type) fed the specific diets 3 days prior to oral gavage. The mice from the same group (4 mice) were housed in the same biocontainment cages (Allentown Inc.) for the duration of the experiment. Mice were maintained on autoclaved water. Fecal samples were collected every 2-3 days after oral gavage for NGS sequencing and pH measurement. The pH of diluted fecal and cecal samples was calculated by measuring the supernatant of 0.1% (v/v) dilution in molecular grade water using a pH probe (Mettler-Toledo)^71^. At the end of the experiment, mice were euthanized, and the cecal contents were collected for NGS sequencing and pH measurement.

### Genomic DNA extraction from fecal and cecal samples

The DNA extraction for fecal and cecal samples was performed as described previously with some modifications^72^. Fecal samples (∼50 mg) were transferred into solvent-resistant screw-cap tubes (Sarstedt Inc) with 500 μL 0.1 mm zirconia/silica beads (BioSpec Products) and one 3.2 mm stainless steel bead (BioSpec Products). The samples were resuspended in 500 μL of Buffer A (200 mM NaCl (DOT Scientific), 20 mM EDTA (Sigma) and 200 mM Tris·HCl pH 8.0 (Research Products International)), 210 μL 20% SDS (Alfa Aesar) and 500 μL phenol/chloroform/isoamyl alcohol (Invitrogen). Cells were lysed by mechanical disruption with a bead-beater (BioSpec Products) for 3 min twice to prevent overheating. Next, cells were centrifuged for 5 min at 8,000 x g at 4°C, and the supernatant was transferred to a Eppendof tube. We added 60 μL 3M sodium acetate (Sigma) and 600 μL isopropanol (LabChem) to the supernatant and incubated on ice for 1 hr. Next, samples were centrifuged for 20 min at 18,000 x g at 4°C. The harvested DNA pellets were washed once with 500 μL of 100% ethanol (Koptec). The remaining trace ethanol in the sample was removed by air drying. Finally, the DNA pellets were then resuspended into 200 μL of AE buffer (Qiagen) left in 4°C overnight to facilitate dissolving. The crude DNA extracts were purified by a Zymo DNA Clean & Concentrator™-5 kit (Zymo Research). The following PCR amplification and NGS sequencing procedures are the same as previously described for treatment of *in vitro* samples.

### GC quantification of organic acids from cecal samples

Organic acids were quantified using headspace gas chromatography (HS-GC). Frozen cecal samples (15 ∼ 40 mg) were prepared for by adding the weighed samples to chilled 20 mL glass vials (Restek, Bellefonte, PA) with 2.0 g of sodium hydrogen sulfate for acidification, distilled water (300 µL – mg of cecal content), and 1.0 mL of 500 µM 2-butanol as an internal standard. Glass vials were immediately sealed with aluminum crimp caps with rubber septa (Restek, Bellefonte, PA), vigorously mixed by hand to mix contents, and left at room temperature overnight. Prepared samples were loaded on a HS-20 headspace sampler (Shimadzu, Columbia, OH) connected to a Shimadzu GC-2010 Plus GC with a flame ionization detector. The column used was an SH-Stabilwax (30 m, 0.25 mm internal diameter, 0.10 µM film thickness). The GC protocol used is described in Hutchison et al. (2023)^73^. Standards for acetate, propionate, isobutyrate, butyrate, and isovalerate, and valerate were combined and serially diluted to generate a standard curve. Valerate was not detected in the cecal samples. Analytes measured had their areas under the curve calculated with Shimadzu Lab Solution (Ver. 5.92) and concentrations were normalized by sample mass and converted to µmol/g using a standard curve.

### Gene expression measurements of the *arc* operon in germ-free mice

The RNA extraction of the contents of each cecum was performed as described previously with some modifications^74^. Specifically, cecal samples (100∼150 mg) were transferred into solvent-resistant screw-cap tubes (Sarstedt Inc) with 500 μL of acid washed beads (212-300 μm, Sigma). The samples were resuspended in 500 μL of Acid-Phenol:Chloroform (with IAA, 125:24:1, pH 4.5, Thermo Fisher Scientific), 500 μL of Buffer B (200mM NaCl (DOT Scientific), 20 mM EDTA (Sigma), pH 8.0), 210 μL 20% SDS (Alfa Aesar). Cells were lysed by mechanical disruption with a bead-beater (BioSpec Products) for 2 min twice to prevent overheating. Next, cells were centrifuged for 10 min at 8,000 x g at 4°C, and the supernatant was transferred into an Eppendof tube. We added 60 μL 3M sodium acetate (pH 5.5, Sigma) and 600 μL isopropanol (LabChem) to the supernatant and incubated at −80°C for 6 min. Next, samples were centrifuged for 15 min at 18,200 x g at 4°C. The harvested RNA pellets were washed once with 900 μL of 100% ethanol (Koptec). Finally, the RNA pellets were resuspended into 100 μL of RNase-free water. The crude RNA extracts were purified by a RNeasy Mini kit (Qiagen) with a DNase I (Qiagen) treatment step to eliminate DNA contamination in the sample. To remove any remaining DNA, the RNA products were treated with Baseline-ZERO DNase I (Epicentre) and then purified again using the RNeasy Mini kit.

We performed cDNA synthesis with 0.5-1 μg of total purified RNA using the iScript Select cDNA Synthesis Kit (Bio-Rad Laboratories). We performed quantitative reverse transcription PCR (qRT-PCR) on the Bio-Rad CFX connect Real-Time PCR instrument with SYBR™ Green PCR Master Mix (Thermo Fisher Scientific) using primers qPCR_arc_fwd and qPCR_arc_rev for arc expression measurement with three biological replicates and 3 technical replicates for each biological replicate. We also included qPCR controls measuring the 16S rRNA level (primers: qPCR_EC16S_fwd and qPCR_EC16S_rev). We computed the fold changes of the target genes by normalizing to the reference gene 16S rRNA gene using the geometric mean through the 2^-ΔΔ*cq*^ method^75^, where:

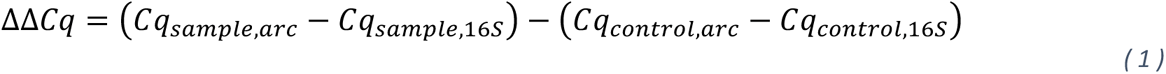

### Bioinformatic analysis for *arc*-harboring species identification

The previous human microbiome metagenome-assembled genome (MAG) dataset containing 154,723 MAGs from 9,428 human gut microbiomes from diverse geographic location, body sites, diseases, and lifestyles was used as the reference database^76^. These genomes extend to > 95% the median mappability of gut microbiomes. All MAGs were annotated by Prodigal v2.6.3^77^. Only genomes with >= 95% completeness and < 10% contamination were used in the downstream analysis (45,502 MAGs). The KOfam HMMs^78^ (KOfam 27-Aug-2021 release) for the four *arc* operon genes (K01478, arcA; K00611, arcB; K00926, arc; K03758, arcD) were used as the reference to search against all the MAGs by using hmmsearch implemented in HMMER v3.1b2^79^. The hmmsearch hits and genome taxonomy were summarized accordingly.

### pH model predictions and anlysis

The pH model fitted on monoculture growth was used to predict community dynamics. Parameters optimized for monoculture of each species of interest were used to simulate community growth curve and pH changes using Python ODE solver scipy.integrate.solve_ivp.

The value of monoculture-pH-divergence for each species in each initial pH condition were calculated by the root mean squared error (RMSE) between model predicted absolute abundance and experimental absolute abundance for each species in each coculture condition averaged across all timepoints, as shown in **equation 7**, where *X_i,pred,t_* represents model predicted absolute abundance of species *i* at timepoint *t* and where *X_i,exp,t_* represents model experimental absolute abundance of species *i* at timepoint *t*. The sign of monoculture-pH-divergence is determined by the sign of *X_i,model,t_* - *X_i,exp,t_*.

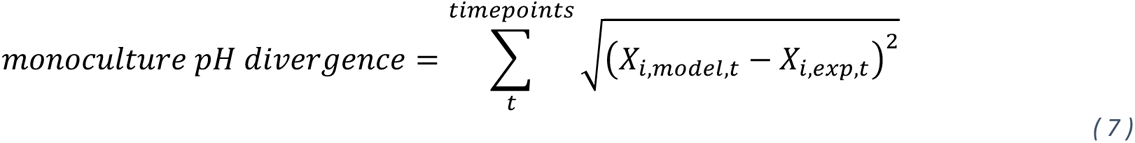

Shannon diversity index for a community was calculated following **equation 8** for both experimental data and model prediction at a given initial pH condition, where *x*_i_ represents relative abundance for species *i*.

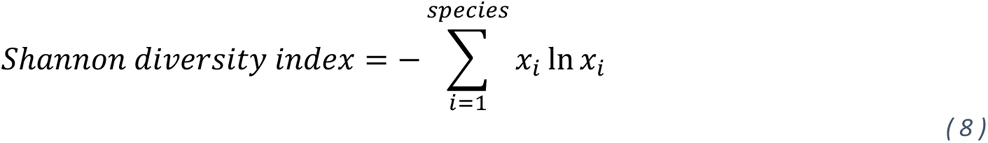

## Data Availability

Additional data supporting the findings described in this work are available from the corresponding author under reasonable request. The processed data for all community experiments and simulation results are available in a Github Repository: https://github.com/VenturelliLab/Liu_et_al_2024.

## Code Availability

Scripts for model training and simulations are available in a Github Repository here: https://github.com/VenturelliLab/Liu_et_al_2024.

## Acknowledgements

We would like to thank all current and past members of the Venturelli lab for their helpful discussions. We would like to thank Ryan Clark, Susan Hromada, and Bryce Connors for their advice related to anaerobic culturing. We would like to thank Jun Feng for their advice on RNA extraction and fecal sample processing techniques. Research was sponsored by the National Institutes of Health under Grant Number R35GM124774, R01EB030340 and Army Research Office W911NF-19-1-0269.

## Author contributions

Y.L, Y.Y.C., and O.S.V conceived the research. Y.L. and Y.Y.C. constructed the plasmids. Y.L. engineered the strains. Y.L. performed the *in vitro* community experiments. Y.L. and J.T. implemented computational modeling. E.I.V. and F.E.R. guided germ-free mouse experiments. M.F.W. performed metabolomics on cecal samples from mice. Z.C.Z. and K.A. performed bioinformatics analysis. O.S.V. secured funding. Y.L. and O.S.V. wrote the manuscript and all authors provided feedback on the manuscript.

## Ethics declarations

The authors declare no competing interests.

