## Supplementary Information for "Shaping human gut community assembly and butyrate production by controlling the arginine dihydrolase pathway"

### Parameter estimation methods

#### Ordinary differential equation model

To model the coupling of species growth with pH over time, we used the following system of ordinary differential equations for each species,

$$\frac{ds}{dt} = s\left(\mu - \frac{s}{K}\right)\left(2^{-\beta(p-p_{\text{pref}})^2} - \delta\right) \tag{1}$$

$$\frac{dp}{dt} = \gamma\frac{ds}{dt}p(1 - p/14) \tag{2}$$

where $s$ is the abundance (OD 600) of species, and $p$ is the pH. The set of model parameters includes the species growth rate $\mu$, carrying capacity $K$, preferred pH, $p_{\text{pref}}$, sensitivity to deviation from the preferred pH, $\beta$, and a constant to relate change in species growth to the change in pH, $\gamma$. For parameters that are strictly nonnegative, such as the growth rate, we define a real-valued parameter as the log of the original parameter, e.g. $\hat{\mu} = \log(\mu)$. The full set of model parameters is denoted as $\theta = \{\log(\mu), \log(K), \log(\beta), \log(p_{\text{pref}}), \log(\delta), \gamma\} \in \mathbb{R}^6$.

#### Parameter estimation

We denote an experimental condition as $q = (s_0, p_0, \tau)$, which specifies the initial condition of species abundance and pH and the measurement time, $\tau$. For each experimental condition, we denote the vector of model predictions of species and pH at time $\tau$ as $\hat{\mathbf{y}}(q, \theta)$, so that

$$\hat{\mathbf{y}}(q, \theta) = \begin{bmatrix} s_0 \\ p_0 \end{bmatrix} + \int_0^{\tau} \begin{bmatrix} s\left(\mu - \frac{s}{K}\right)\left(2^{-\beta(p-p_{\text{pref}})^2} - \delta\right) \\ \gamma\frac{ds}{dt}p(1 - p/14) \end{bmatrix} dt \tag{3}$$

Measurements of species and pH are assumed to be corrupted by zero-mean Gaussian noise, $\mathbf{y}(q) = \hat{\mathbf{y}}(q, \theta) + \varepsilon$, where $\varepsilon \sim \mathcal{N}(0, \nu^{-1}\mathbf{I})$ and $\nu$ is the measurement precision. Given a dataset of measurements for $n$ conditions, $\mathcal{D} = \{\mathbf{y}(q_i), ..., \mathbf{y}(q_n)\}$, the posterior parameter distribution is proportional to the product of data likelihood and a parameter prior distribution,

$$p(\theta|\mathcal{D}, \alpha, \nu) \propto \prod_{i=1}^{n} \mathcal{N}(\mathbf{y}(q_i)|\hat{\mathbf{y}}(q_i, \theta), \nu^{-1}\mathbf{I})\mathcal{N}(\theta|0, \alpha^{-1}\mathbf{I}). \tag{4}$$

where $\alpha$ is the precision of the parameter prior. The *maximum a posteriori* (MAP) parameter estimates are determined by minimizing the negative log of the parameter posterior distribution with respect to $\theta$,

$$\theta_{MAP} \in \underset{\theta}{\operatorname{argmin}} \sum_{i=1}^{n} (\mathbf{y}(q_i) - \hat{\mathbf{y}}(q_i, \theta))^T(\mathbf{y}(q_i) + \hat{\mathbf{y}}(q_i, \theta)) + \lambda \cdot \theta^T\theta \tag{5}$$

where $\lambda = \alpha/\nu$ is the regularization coefficient. The regularization coefficient is a hyperparameter that reduces the propensity of the model to overfit to noise in the data by promoting the emphasis on the prior. Optimization of Eq. 5 is performed using Scipy's MINIMIZE function using the Newton-CG method [4], where the Hessian of the negative log posterior with respect to $\theta$ is computed using the outer product approximation,

$$\mathbf{H} \approx \lambda \mathbf{I} + \sum_{i=1}^{n} \nabla_\theta \hat{\mathbf{y}}(q_i, \theta) \cdot \nabla_\theta \hat{\mathbf{y}}(q_i, \theta)^T. \tag{6}$$

Numerical integration and computation of gradients is performed using the JAX package in Python [2]. Optimization of model parameters is performed 3 times using different initial parameter guesses, where the parameters that resulted in the lowest value for the objective Eq. 5 is kept. Using the optimized parameters as an initial estimate for the mean of the parameter posterior, we use a variational inference [1] algorithm to optimize a Gaussian approximation of the parameter posterior distribution.

#### Variational inference

We use a variational inference approach to approximate the parameter posterior distribution as an independent Gaussian for each parameter, $p(\theta_i|\mathcal{D}) \approx \mathcal{N}(\theta_i|\mu_i, \sigma_i^2)$. Optimization of the approximate parameter posterior, denoted as $q(\theta|\mu, \sigma^2)$, involves minimizing its Kullback-Leibler (KL) divergence with the true posterior,

$$\mu^*, \sigma^* \in \underset{\mu,\sigma}{\operatorname{argmin}} \, \mathrm{KL}\left[q(\theta|\mu,\sigma)||p(\theta|\mathcal{D},\alpha,\nu)\right]. \tag{7}$$

The KL divergence is

$$\begin{aligned} \mathrm{KL}\left[q(\theta|\mu,\sigma)||p(\theta|\mathcal{D},\alpha,\nu)\right] = & \int_\theta q(\theta|\mu,\sigma) \log \frac{p(\theta|\mathcal{D},\alpha,\nu)}{q(\theta|\mu,\sigma)} d\theta \\ = & \int_\theta q(\theta|\mu,\sigma) \log p(\mathcal{D}|\theta,\nu) d\theta + \int_\theta q(\theta|\mu,\sigma) \log p(\theta|\alpha) d\theta \\ & - \int_\theta q(\theta|\mu,\sigma) \log p(\mathcal{D}|\alpha,\nu) d\theta - \int_\theta q(\theta|\mu,\sigma) \log q(\theta|\mu,\sigma) d\theta. \end{aligned} \tag{8}$$

Keeping only the terms that depend on the variational parameters $\mu$ and $\sigma$, we define

$$\mathcal{L}(\mu,\sigma) = \int_\theta q(\theta|\mu,\sigma) \log p(\mathcal{D}|\theta,\nu) d\theta + \int_\theta q(\theta|\mu,\sigma) \log p(\theta|\alpha) d\theta - \int_\theta q(\theta|\mu,\sigma) \log q(\theta|\mu,\sigma) d\theta. \tag{9}$$

The function $\mathcal{L}(\mu,\sigma)$ is called the evidence lower bound (ELBO). Although the expectation of the log likelihood is analytically intractable, the ELBO can be estimated using a Monte Carlo approximation,

$$\mathcal{L}(\mu,\sigma) \approx -\frac{1}{m} \sum_{j=1}^{m} \sum_{i=1}^{n} \sum_{k=1}^{n_y} \nu_k (y_k(q_i) - \hat{y_k}(q_i, \theta^j))^2/2 - \frac{1}{m} \sum_{j=1}^{m} \sum_{l=1}^{n_\theta} \alpha_l (\theta_l^j)^2/2 + \sum_{i=1}^{n_\theta} \log \sigma_i. \tag{10}$$

where $\theta^j \sim \mathcal{N}(\theta|\mu,\sigma)$. Optimization of the ELBO with respect to $\mu$ and $\sigma$ is performed using the ADAM algorithm [3].

#### Expectation Maximization to optimize hyperparameters

Model hyperparameters include the precision of the parameter prior, $\alpha$, and the precision of the measurement noise, $\nu$, which determine the balance of emphasis on the prior and likelihood functions, respectively. The expectation maximization (EM) algorithm [1] iterates between inference of the parameter posterior distribution and then maximzation of the expected joint distribution of the data and model parameters with respect to model hyperparameters. Because inference of the posterior depends on the hyperparameters, the posterior distribution needs to be updated given the optimized hyperparameters. The process of iterating between updating the posterior and updating hyperparameters continues until convergence of the ELBO. The expectation of the joint distribution of the data and model parameters is

$$\begin{aligned}
\mathcal{Q}(\alpha,\nu) &= \int_\theta q(\theta|\mu,\sigma)\log p(\mathcal{D},\theta|\alpha,\nu)d\theta \\
&= \int_\theta q(\theta|\mu,\sigma)\log p(\mathcal{D}|\theta,\nu)d\theta + \int_\theta q(\theta|\mu,\sigma)\log p(\theta|\alpha)d\theta \\
&\approx -\frac{1}{m}\sum_{j=1}^{m}\sum_{i=1}^{n}\frac{1}{2}\left(\sum_{k=1}^{n_y}\nu_k(y_k(q_i)-\hat{y_k}(q_i,\theta^j))^2-\log\nu_k\right)-\frac{1}{m}\sum_{j=1}^{m}\frac{1}{2}\sum_{l=1}^{n_\theta}\left(\alpha_l(\theta_l^j)^2-\log\alpha_l\right)
\end{aligned} \tag{11}$$

where $\theta^j \sim \mathcal{N}(\theta|\mu,\sigma)$. Maximization of $\mathcal{Q}(\alpha,\nu)$ with respect to $\alpha$ and $\nu$ gives the update equations,

$$\alpha_l = \frac{1}{\frac{1}{m}\sum_{j=1}^{m}(\theta_l^j)^2} \tag{12}$$

$$\nu_k = \frac{1}{m}\sum_{j=1}^{m}\frac{1}{n}\sum_{i=1}^{n}\frac{1}{(y_k(q_i)-\hat{y_k}(q_i,\theta^j))^2} \tag{13}$$

for $l \in \{1,...,6\}$ and $k \in \{1,2\}$.

[4] Pauli Virtanen, Ralf Gommers, Travis E. Oliphant, Matt Haberland, Tyler Reddy, David Cournapeau, Evgeni Burovski, Pearu Peterson, Warren Weckesser, Jonathan Bright, Stéfan J. van der Walt, Matthew Brett, Joshua Wilson, K. Jarrod Millman, Nikolay Mayorov, Andrew R. J. Nelson, Eric Jones, Robert Kern, Eric Larson, C J Carey, İlhan Polat, Yu Feng, Eric W. Moore, Jake VanderPlas, Denis Laxalde, Josef Perktold, Robert Cimrman, Ian Henriksen, E. A. Quintero, Charles R. Harris, Anne M. Archibald, Antônio H. Ribeiro, Fabian Pedregosa, Paul van Mulbregt, and SciPy 1.0 Contributors. SciPy 1.0: Fundamental Algorithms for Scientific Computing in Python. *Nature Methods*, 17:261–272, 2020.

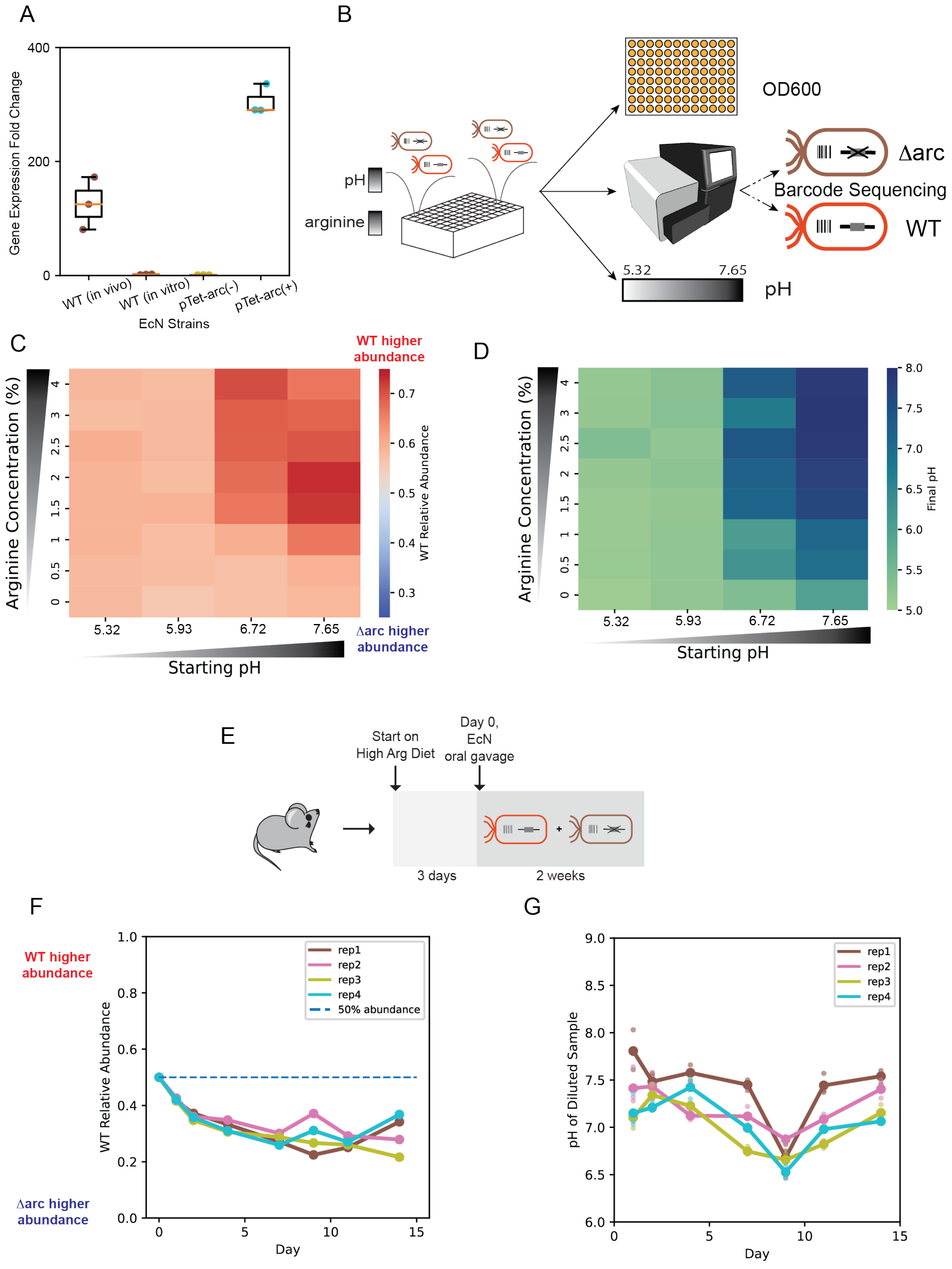

A
Gene Expression Fold Change
400
200
0
WT (in vivo)
WT (in vitro)
pTet-arc(-)
pTet-arc(+)
EcN Strains
B
OD600
pH
arginine
Δarc
Barcode Sequencing
WT
5.32
7.65
pH
C
WT higher abundance
Arginine Concentration (%)
4
3
2.5
2
1.5
1
0.5
0
5.32
5.93
6.72
7.65
Starting pH
0.7
0.6
0.5
0.4
0.3
WT Relative Abundance
Δarc higher abundance
D
Arginine Concentration (%)
Starting pH
8.0
7.5
7.0
6.5
6.0
5.5
5.0
Final pH
E
Start on High Arg Diet
Day 0, EcN oral gavage
3 days
2 weeks
F
WT higher abundance
Δarc higher abundance
WT Relative Abundance
1.0
0.8
0.6
0.4
0.2
0.0
0
5
10
15
Day
rep1
rep2
rep3
rep4
50% abundance
G
pH of Diluted Sample
9.0
8.5
8.0
7.5
7.0
6.5
6.0
0
5
10
15
Day
rep1
rep2
rep3
rep4

**Supplementary Figure 1**. ***E. coli*** **Nissle fitness in mammalian gut is impacted by the *arc* operon.** (A) Box plot of transcriptional fold changes using qRT-PCR of representative genes in the *arc* operon in germ free mice colonized with community and WT strain of EcN compared to *in vitro* monoculture of EcN strains (WT, pTet-arc (-), pTet-arc (+)) in media with 1% arginine supplement. The pTet-arc(-) condition represents pTet-arc strain grown in absence of inducer aTc whereas the pTet-arc(+) condition represents pTet-arc strain grown in presence of inducer aTc. The *in vitro* expression of *arc* operon in pTet-arc (-) was selected as control. Data points represent mean fold change for each biological replicate with three technical replicates for each gene. The distribution of biological replicates in each condition is represented by box plot. Color of data points represent different EcN strains. (B) Experimental workflow of *in vitro* competition experiment between WT and Δarc strains. Strains were barcoded and cocultured in arginine and initial media pH titrations. Batch cultures were grown for 24 hours at 37°C until harvesting for measurements of OD600, supernatant pH, and barcode sequencing. (C) Heatmap of WT strain relative abundance in coculture under arginine and initial media pH titration. (D) Heatmap of supernatant pH after 24 hours of coculture under arginine and initial media pH titration. (E) Experimental workflow of *in vivo* competition experiment between WT and Δarc strains. Mice were tagged and placed on customized high arginine diet 3 days before oral gavage of equal-part barcoded WT and Δarc strains. During two weeks of co-colonization, fecal samples were collected every two to three days for barcode sequencing and pH measurement. (F) Relative abundance of WT strain in fecal and cecal sample, with replicates shown in different color. The equal colonization is marked by blue dashed line at 50% abundance (G) The pH of diluted fecal and cecal samples, with replicates shown in different color.

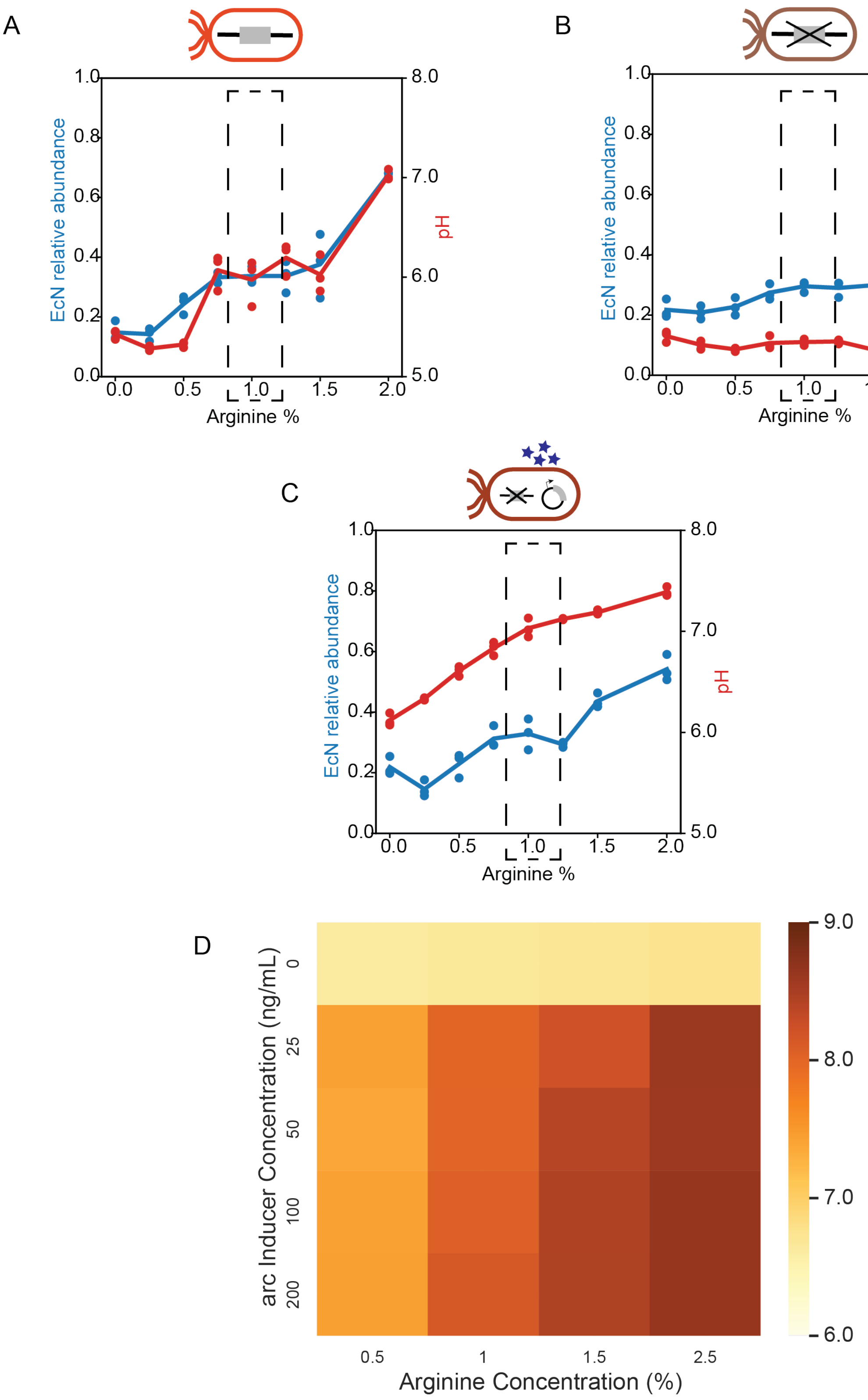


**Supplementary Figure 2**. **Relationship between arginine, EcN abundance and endpoint pH.** (A), (B), and (C) EcN relative abundance and final pH of community coculture with WT (A), Δarc

(B), and pTet-arc (+) (C) strains respectively. EcN relative abundances are represented by blue and final pHs are represented by red. EcN strains and community species were precultured in YBHI media for 16 hours before diluting to 0.01 initial OD600 into DM38 for each species. EcN relative abundance is calculated based on 16S NGS sequencing result, dividing EcN read counts by total read counts. Final pH is measured by the PhenolRed method. Individual replicates are shown as circles and average of three replicates is shown as line. Dashed rectangle indicate the experimental condition selected for this study. (D) Final pH of pTet-arc (+) batch culture with arginine and aTc inducer concentration titration. EcN strain pTet-arc (+) was precultured in YBHI before diluting to 0.01 OD600 into fresh DM38 media with the arginine and inducer concentrations indicated in the figure. Dashed rectangle indicates the experimental condition selected for this study. Each condition contained 3 biological replicates.

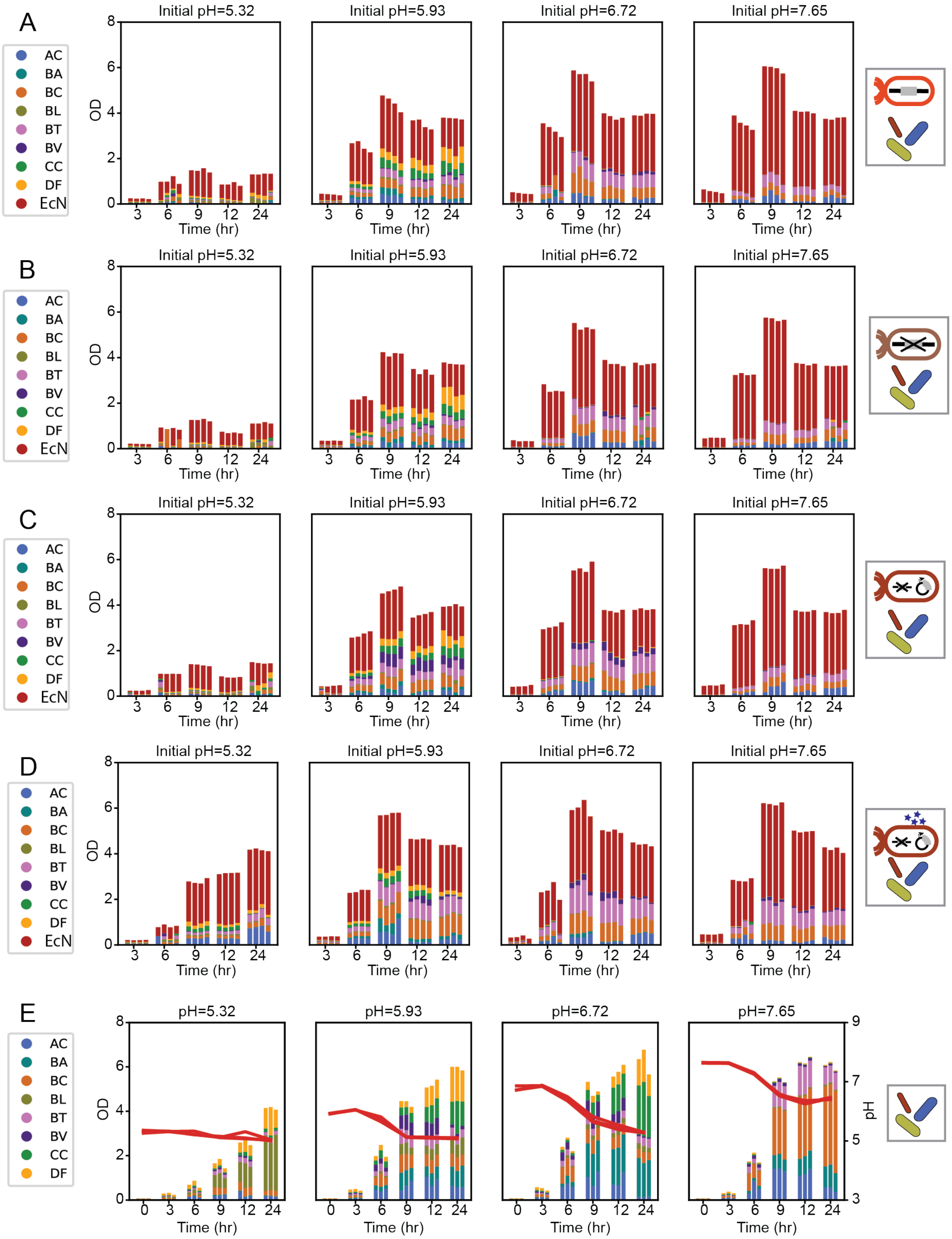


**Supplementary Figure 3**. **Species absolute abundance across multiple timepoints *in vitro*.** (A), (B), (C), and (D) Species absolute abundance bar plot of *in vitro* coculture of community with WT (A), Δarc (B), pTet-arc (-) (C), and pTet-arc (+) (D) strains, respectively. EcN strains and community species were precultured in YBHI media for 16 hours before diluting to 0.01 initial

OD600 into DM38 for each species. Absolute abundances for each species are calculated based on 16S NGS sequencing result, dividing species read counts by total read counts and multiplying the OD600 measurement of a given well. (E) Species absolute abundance bar plot of *in vitro* coculture of community. Community species were precultured in YBHI media for 16 hours before diluting to 0.01 initial OD600 into DM38 for each species. Absolute abundances for each species are calculated based on 16S NGS sequencing result, dividing species read counts by total read counts and multiplying the OD600 measurement of a given well. Timepoint pH measurements were obtained with the PhenolRed method. All three replicates are shown in the bar plots for a given condition.

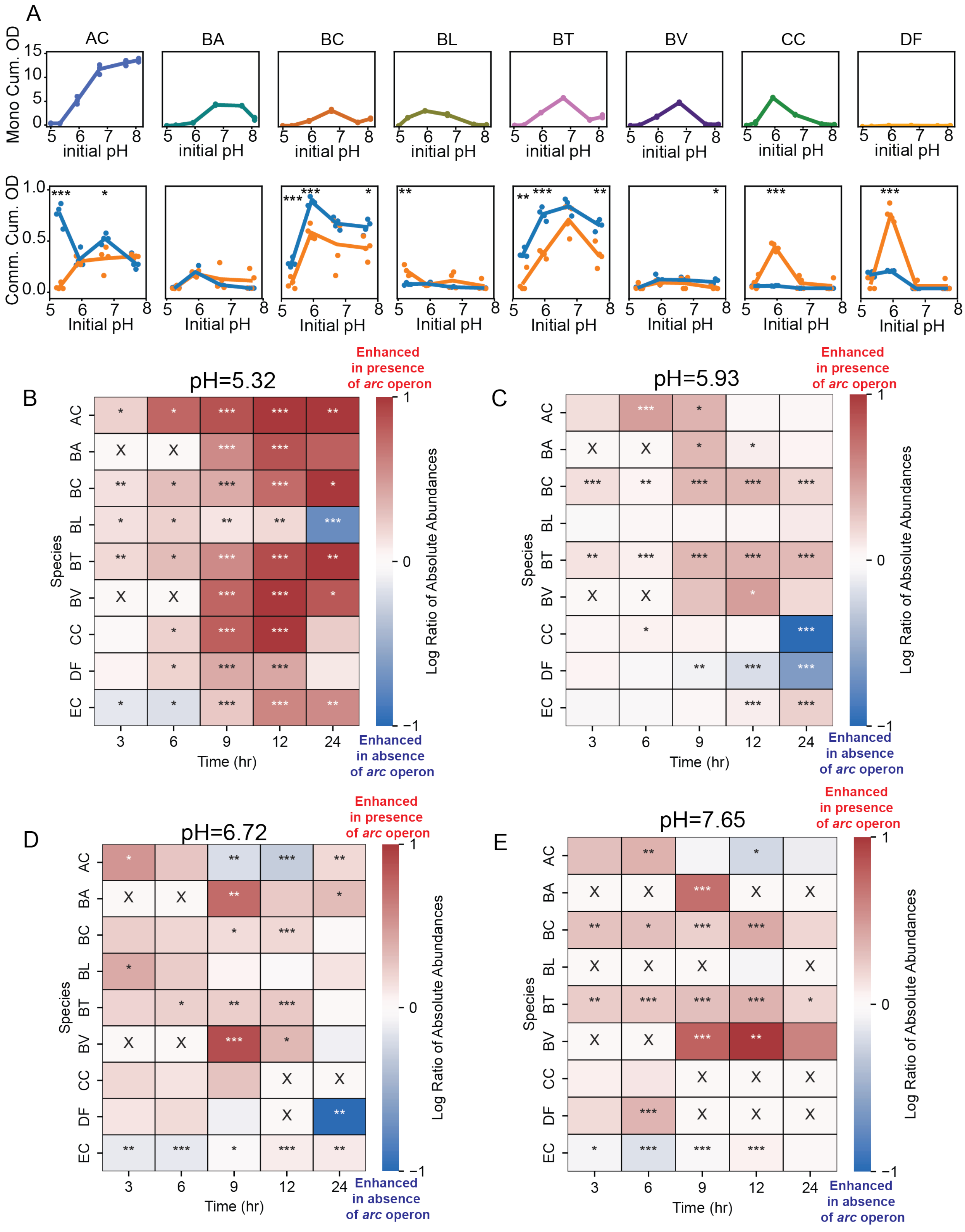
A
AC
BA
BC
BL
BT
BV
CC
DF
Mono Cum. OD
initial pH
Comm. Cum. OD
Initial pH
B
pH=5.32
C
pH=5.93
D
pH=6.72
E
pH=7.65
Species
AC
BA
BC
BL
BT
BV
CC
DF
EC
Time (hr)
3
6
9
12
24
Log Ratio of Absolute Abundances
Enhanced in presence of arc operon
Enhanced in absence of arc operon

**Supplementary Figure 4**. **Changes in species abundance in the presence and absence of the *arc* operon.** (A) Species integral OD of monoculture (top) at each initial pH and species integral absolute abundance in community (bottom) with pTet-arc (+) condition (blue) and Δarc condition (orange) at each initial pH. The species integral OD was calculated by summing OD measured at each timepoint. The species integral absolute abundance in community was calculated by summing absolute abundance calculated at each timepoint of community coculture. The absolute abundance in community at each timepoint was determined by community OD multiplying species relative abundance calculated from NGS data. Color represents species identity. Line represents average of biological replicates (n=4). (B), (C), (D), and (E) Heatmap of the log of the ratio between species absolute abundance (mean value of biological replicates, n = 4) in pTet-arc (+) condition and Δarc condition at initial pH 5.32, 5.93, 6.72, and 7.65, respectively. Species relative abundances are determined by multiplexed 16S rRNA sequencing. Red corresponds to higher relative abundance in pTet-arc (+) conditions and blue corresponds to higher relative abundance in Δarc conditions. Low growth conditions are marked by an 'X'. Asterisks represent statistical significance: $^{*}P < 0.05$, $^{**}P < 0.01$, $^{***}P < 0.001$ according to an unpaired t-test.

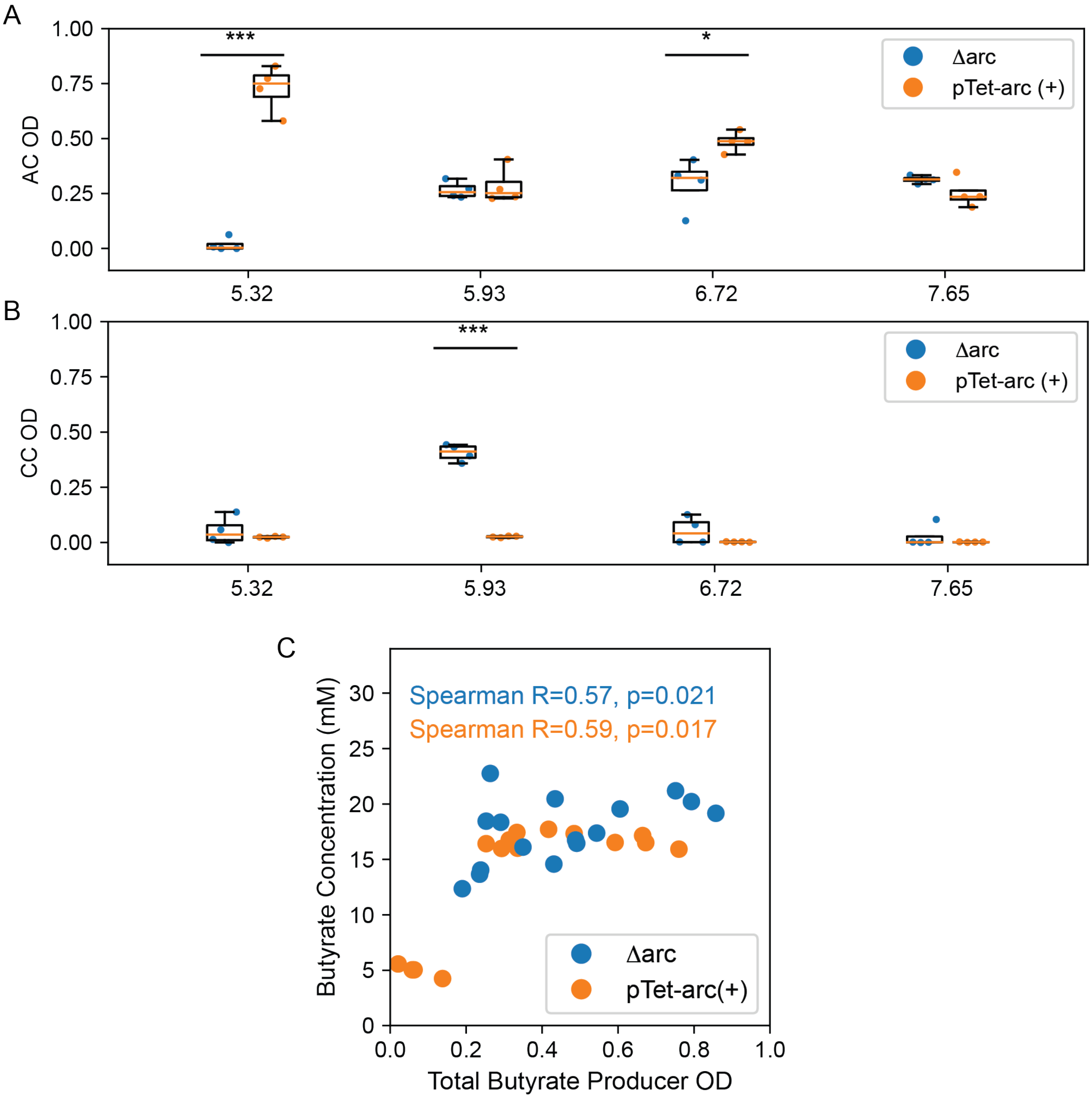


**Supplementary Figure 5**. **Butyrate producer abundances and correlation with butyrate concentration.** (A), (B) The calculated species OD of butyrate producing species AC and CC, respectively. The species OD was calculated by multiplying coculture OD with species relative abundance obtained from NGS. Datapoints indicate experimental data replicates. Datapoint colors indicate strain of EcN in coculture. (C) Scatterplot of final butyrate concentration against total butyrate producer OD at various initial pH. Datapoints indicate experimental data replicates in various initial pH. Datapoint colors indicate strain of EcN in coculture. Spearman correlation is calculated for each strain condition, denoted by color.

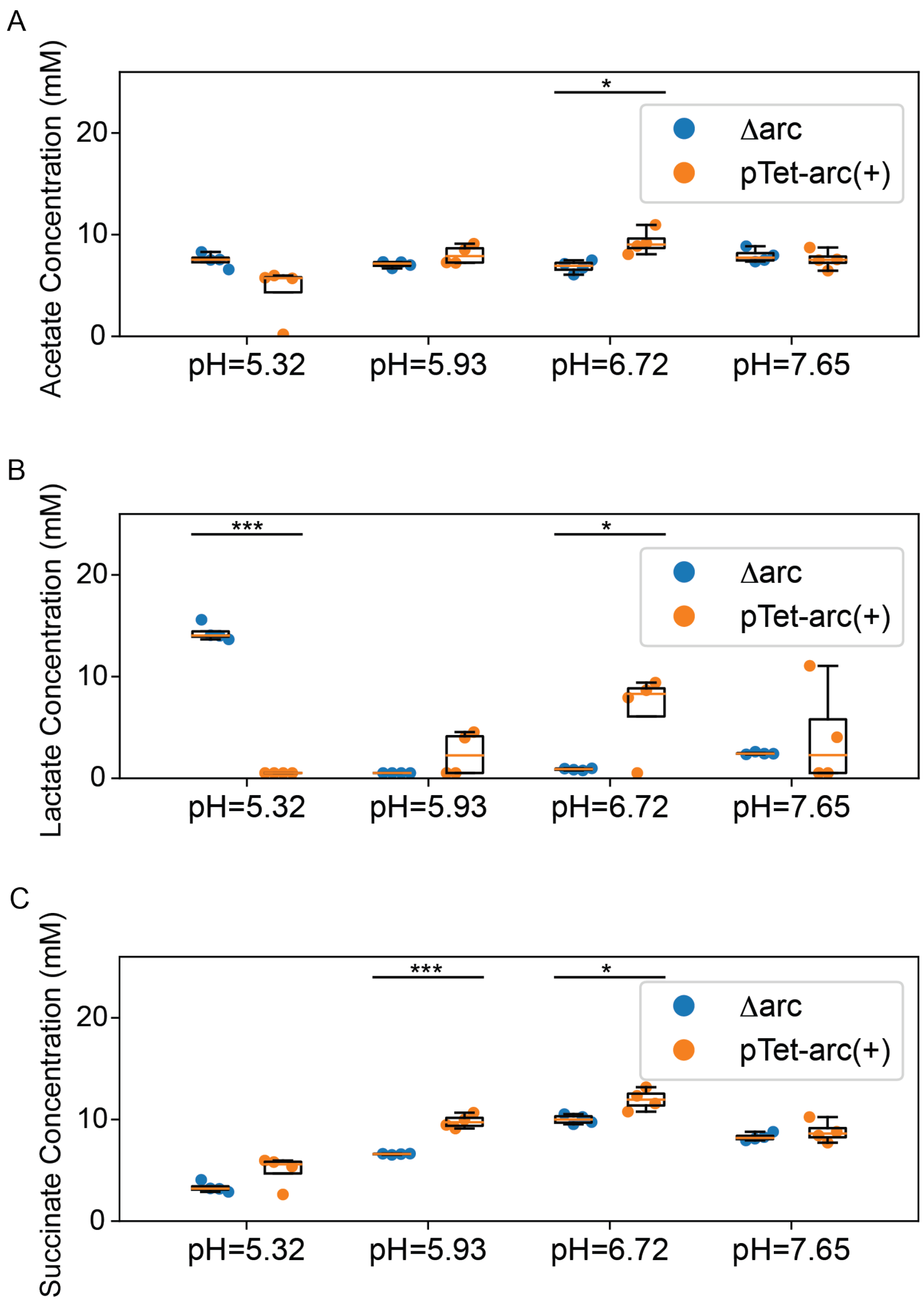


**Supplementary Figure 6. Fermentation end product measurements at 24-hour in human gut communities.** The acetate (A), lactate (B), and succinate (C) concentrations (mM) in supernatant were measured for community cocultures with Δarc (blue) and pTet-arc(+) (orange) strains of EcN at 24 hour by HPLC. Datapoints indicate experimental data replicates. Datapoint colors indicate strain of EcN in coculture. Independent statistical t-test is performed on each pair of conditions with the same starting pH. Asterisks represent statistical significance: *$P < 0.05$, **$P < 0.01$, ***$P < 0.001$ according to an unpaired t-test.

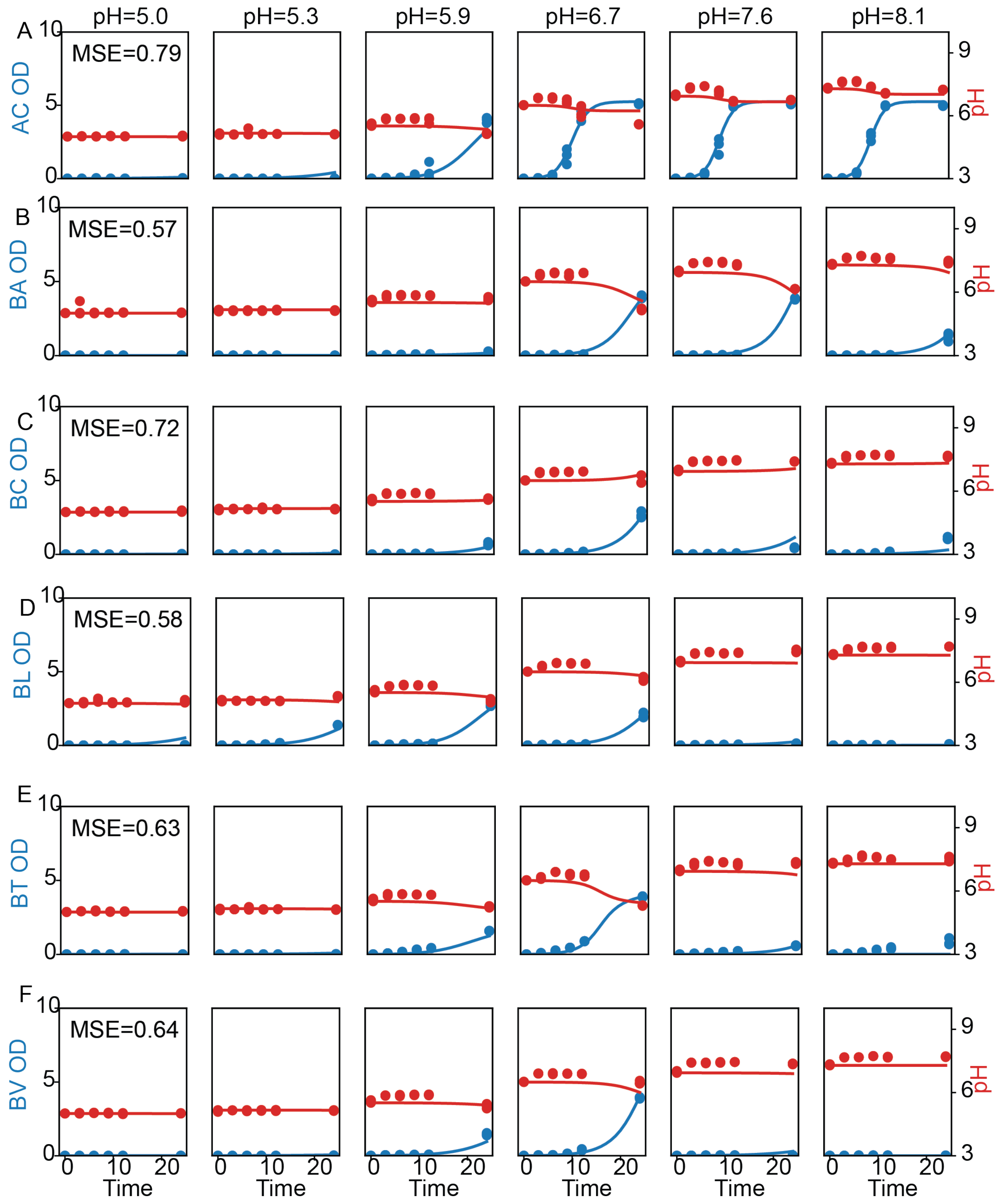


**Supplementary Figure 7**. **pH computational model fit to growth dynamics of AC, BA, BC, BL, BT, and BV.** Plots are shown for AC (A), BA (B), BC (C), BL (D), BT (E), and BV (F). OD data is represented by blue and pH data is represented by red. Experimental data are shown in circles with 3 replicates and model fittings are shown in lines. The mean squared error (MSE) of fitting for each species is displayed on graph, calculated by the mean squared difference between experimental and prediction of OD and pH across all timepoints.

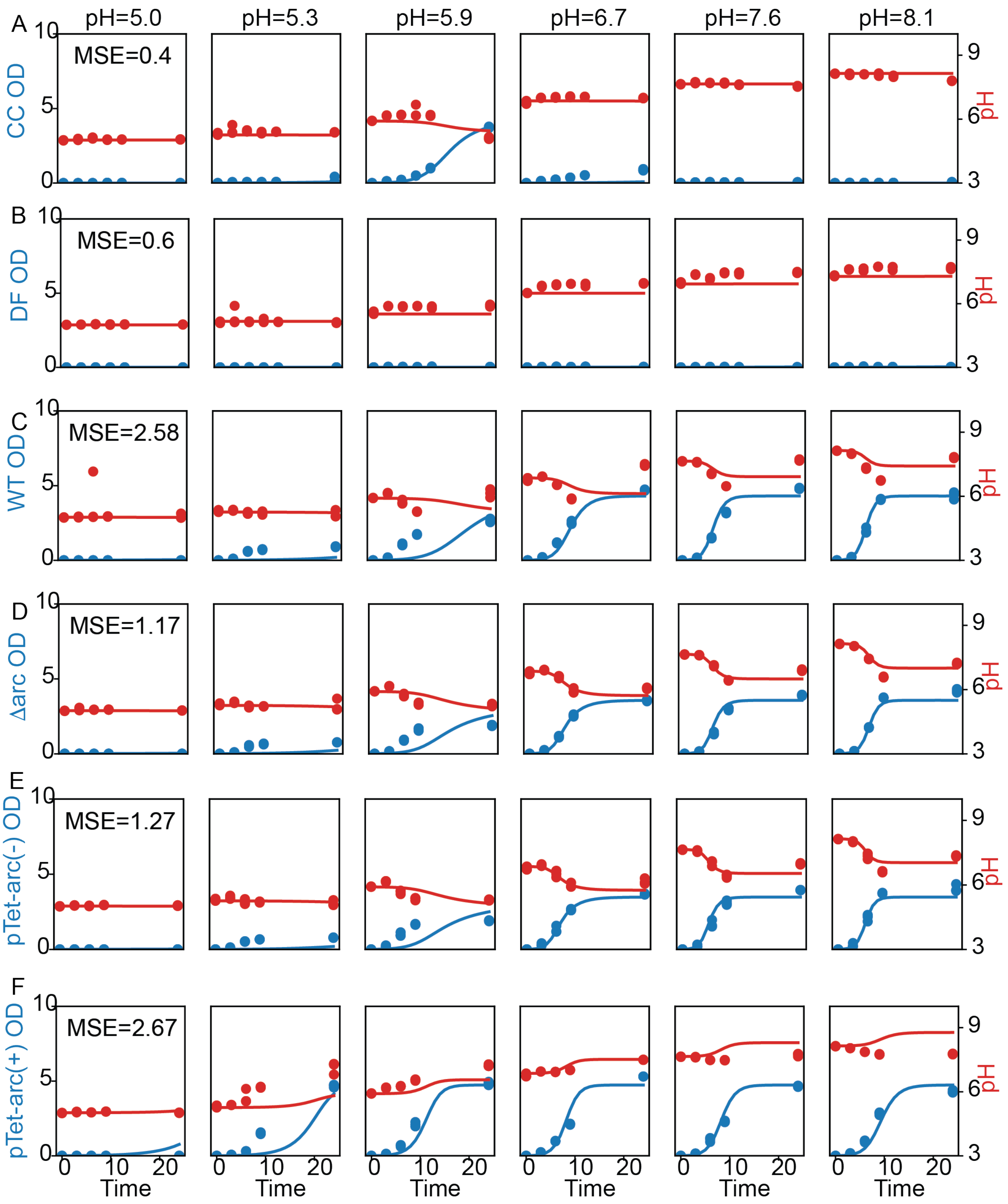


**Supplementary Figure 8**. **pH computational model fit to monoculture dynamics of CC, DF, WT, Δarc, pTet-arc (-), and pTet-arc(+).** Plots are shown for CC (A), DF (B), WT (C), Δarc (D), pTet-arc (E), and pTet-arc (+) (F). OD data is represented by blue and pH data is represented by red. Experimental data are shown in circles with 3 replicates and model fittings are shown in lines. The mean squared error (MSE) is calculated between the experimental and model prediction of OD and pH across all timepoints.

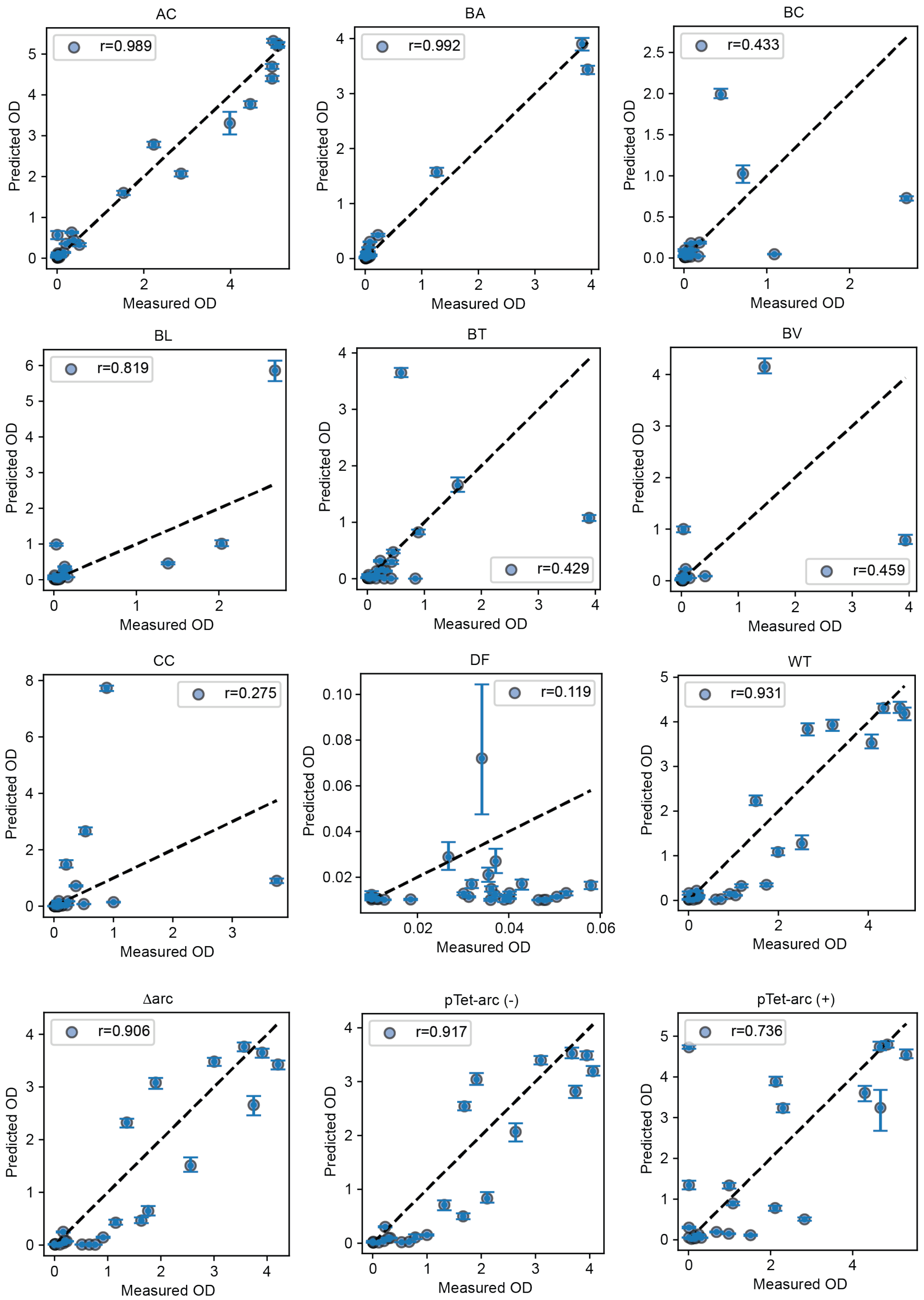

AC
r=0.989
BA
r=0.992
BC
r=0.433
BL
r=0.819
BT
r=0.429
BV
r=0.459
CC
r=0.275
DF
r=0.119
WT
r=0.931
Δarc
r=0.906
pTet-arc (-)
r=0.917
pTet-arc (+)
r=0.736
Predicted OD
Measured OD

**Supplementary Figure 9**. **Prediction performance for each species from leave-one-out cross-validation.** For each species, measurements of species OD and pH over a 24-time interval from 5 of the 6 pH conditions was used for training and the remaining pH condition was held out for testing. Prediction performance was evaluated using the Pearson correlation coefficient between measured and predicted species OD.

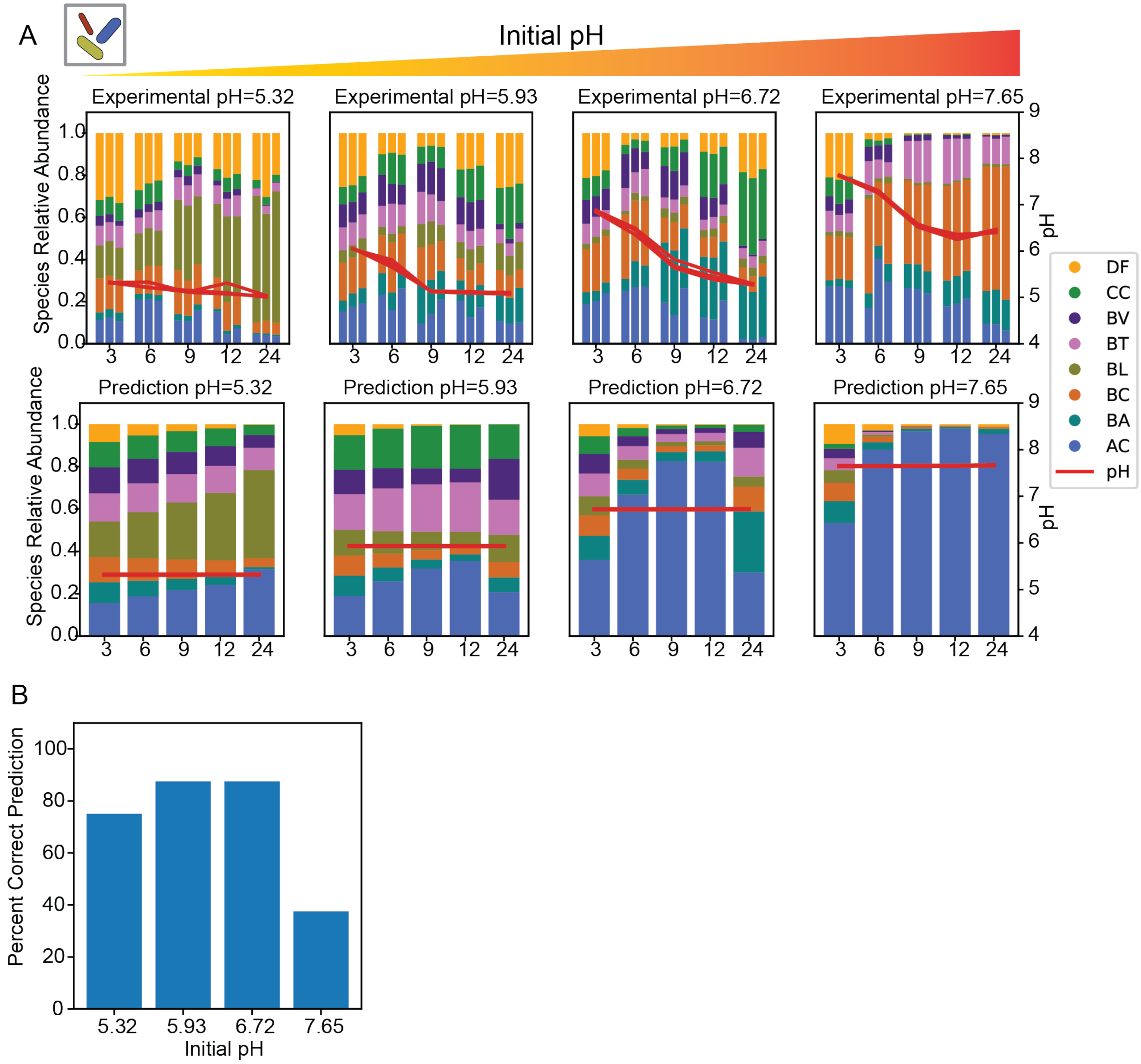


**Supplementary Figure 10. Experimental data and model prediction of human gut community.** (A) Bar plot of experimental composition, predicted composition, and mean squared error of prediction of the 8-member community at various initial pH. Species relative abundances are determined by multiplexed 16S rRNA sequencing. Color indicates species identity. The red line indicates pH measured by PhenolRed method as described in the method section. Measurements and predictions are taken at 3, 6, 9, 12, and 24 hr. Individual biological replicates are shown as individual bars. (B) Bar plot of percent of model correct prediction on species presence/absence for 8-member community in various initial pH. Percentage represents model prediction correctness on absolute abundance where threshold is set at 0.05 (5 times initial inoculum OD of 0.01) for presence and absence of a particular species. Prediction was made for all biological replicates of community assembly measured at 24 hr by 16S sequencing.

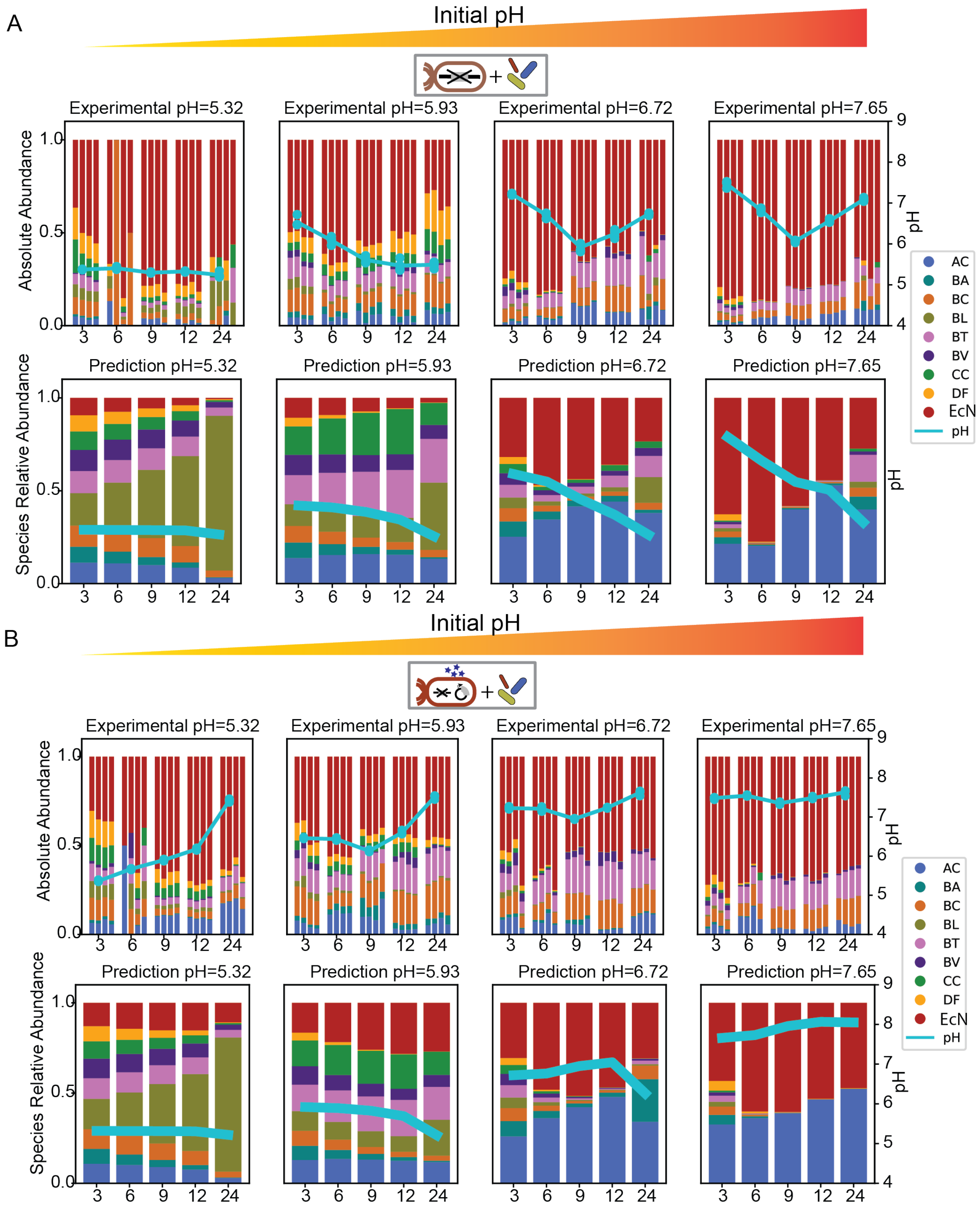


**Supplementary Figure 11. Experimental data and pH computational model predictions of the human gut community containing different EcN strains.** (A) Bar plot of experimental and predicted composition of community coculture with Δarc. Color indicates species identity. The cyan line indicates pH. (B) Bar plot of experimental and predicted composition of community coculture with pTet-arc (+). Color indicates species identity. The cyan line indicates pH.

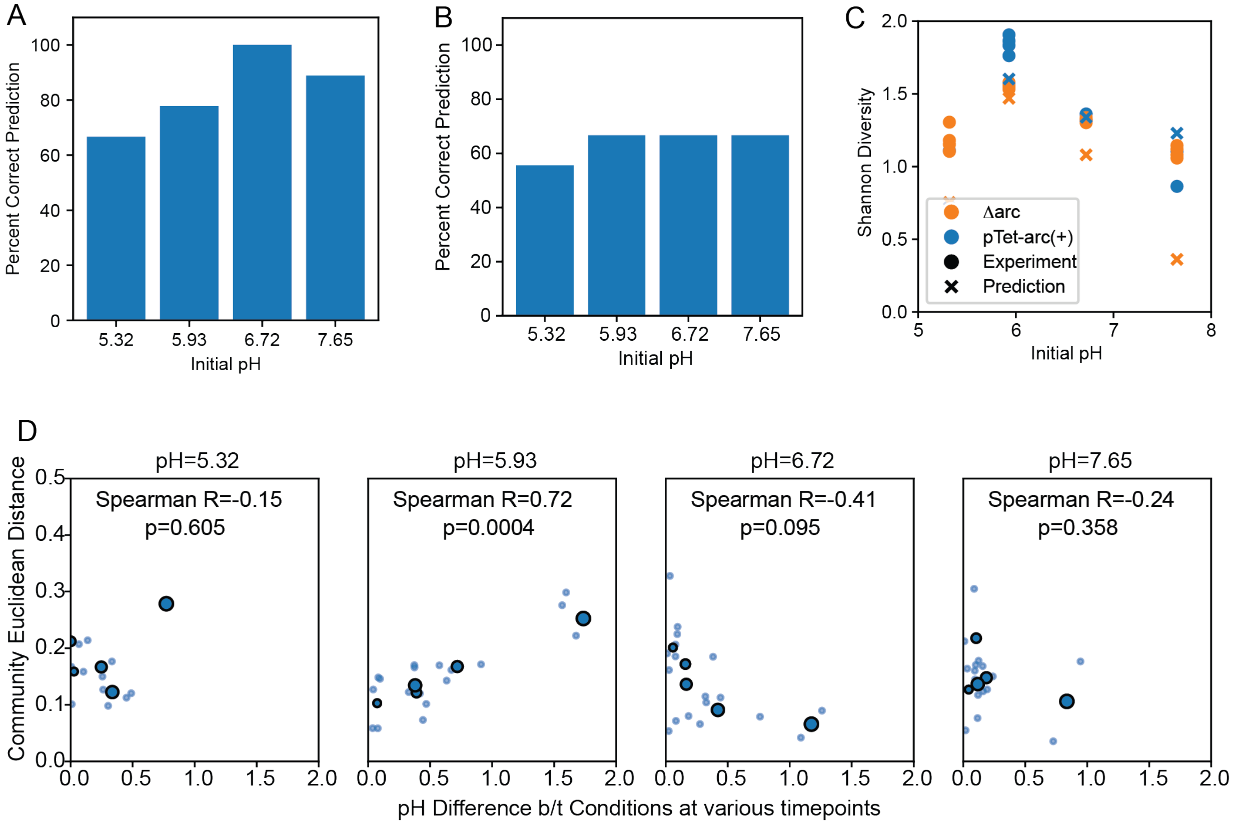


**Supplementary Figure 12**. **Model prediction performance of the presence/absence of species, Shannon diversity and relationship between changes in pH and changes in species abundance.** (A) Bar plot of percent of model correct prediction on species presence/absence for community+Δarc in various initial pH. Percentage represents model prediction correctness on absolute abundance greater than 0.05 (presence) or less than 0.05 (absence). Threshold is set to be 5 times initial inoculum OD at 0.01. Prediction was made for all biological replicates of community assembly measured at 24 hr by 16S sequencing. (B) Bar plot of percent of model correct prediction on species presence/absence for community+pTet-arc(+) in various initial pH. Percentage represents model prediction correctness on absolute abundance greater than 0.05 (presence) or less than 0.05 (absence). Threshold is set to be 5 times initial inoculum OD at 0.01. Prediction was made for all biological replicates of community assembly measured at 24 hr by 16S sequencing. (C) Shannon diversity index of experiment and model prediction of cocultures with pTet-arc(+) and cocultures with Δarc at 24 hr timepoints. Shannon diversity index is calculated for community structures at an initial pH at 24 hr, with equations detailed in the method section. Datapoints represent experimental data and crosses represent model prediction. Colors represent coculture condition with either Δarc (orange) or pTet-arc(+) (blue). (D) Coculture community structure Euclidean distance versus pH difference at various timepoints between community cocultures with pTet-arc (+) and cocultures with Δarc with initial pH at 5.32, 5.93, 6.72, and 7.65, respectively. The Euclidean distance is calculated for community structures at a given timepoint between pTet-arc (+) conditions and Δarc conditions. Timepoint pH values are measured by PhenolRed method as described in the method section. The pH differences are calculated between pH measurements made for different coculture condition at the same timepoints. Solid blue data points represent individual timepoints at an initial pH and light blue datapoints represent 4 biological replicates. Solid blue datapoint size represent timepoints. Spearman $\rho$ and p-value are shown as text in each subplot.

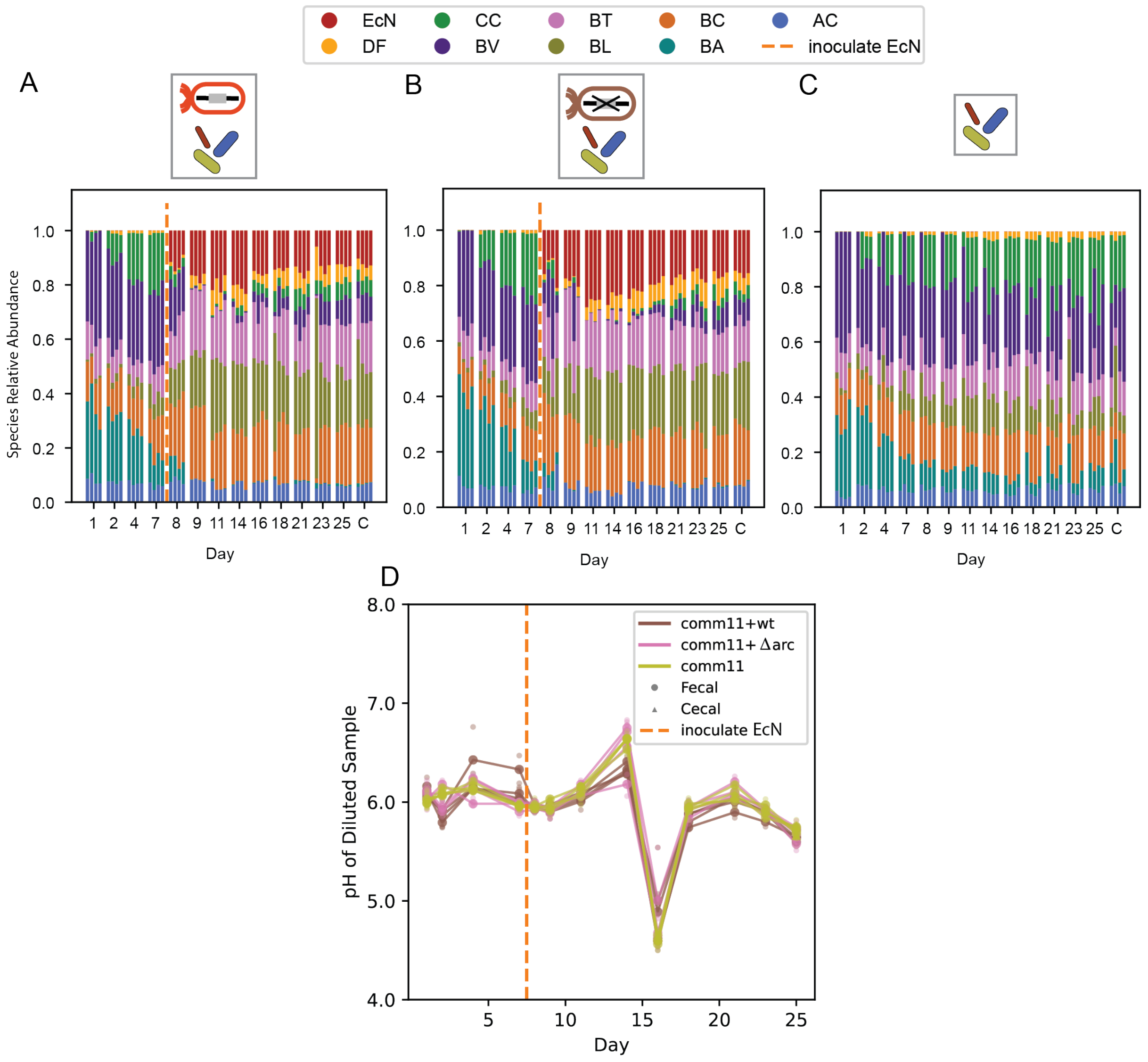


**Supplementary Figure 13**. **Species relative abundance and pH of fecal/cecal sample across multiple timepoints in mice.** (A), (B), and (C) Species relative abundance bar plot of *in vivo* co-colonization of community with WT (A), Δarc (B), and no EcN strains (C), respectively. Species of community were introduced to mice by oral gavage at equal initial OD600. Color indicates species identity. All four replicates are shown for each experimental condition. (D) The pH of diluted fecal and cecal samples. The diluted sample pH is measured by pH meter. Datapoints indicate experimental data replicates. Lines indicate average over all biological replicates. Line colors indicate different experimental condition. EcN inoculation is marked by orange dashed line.

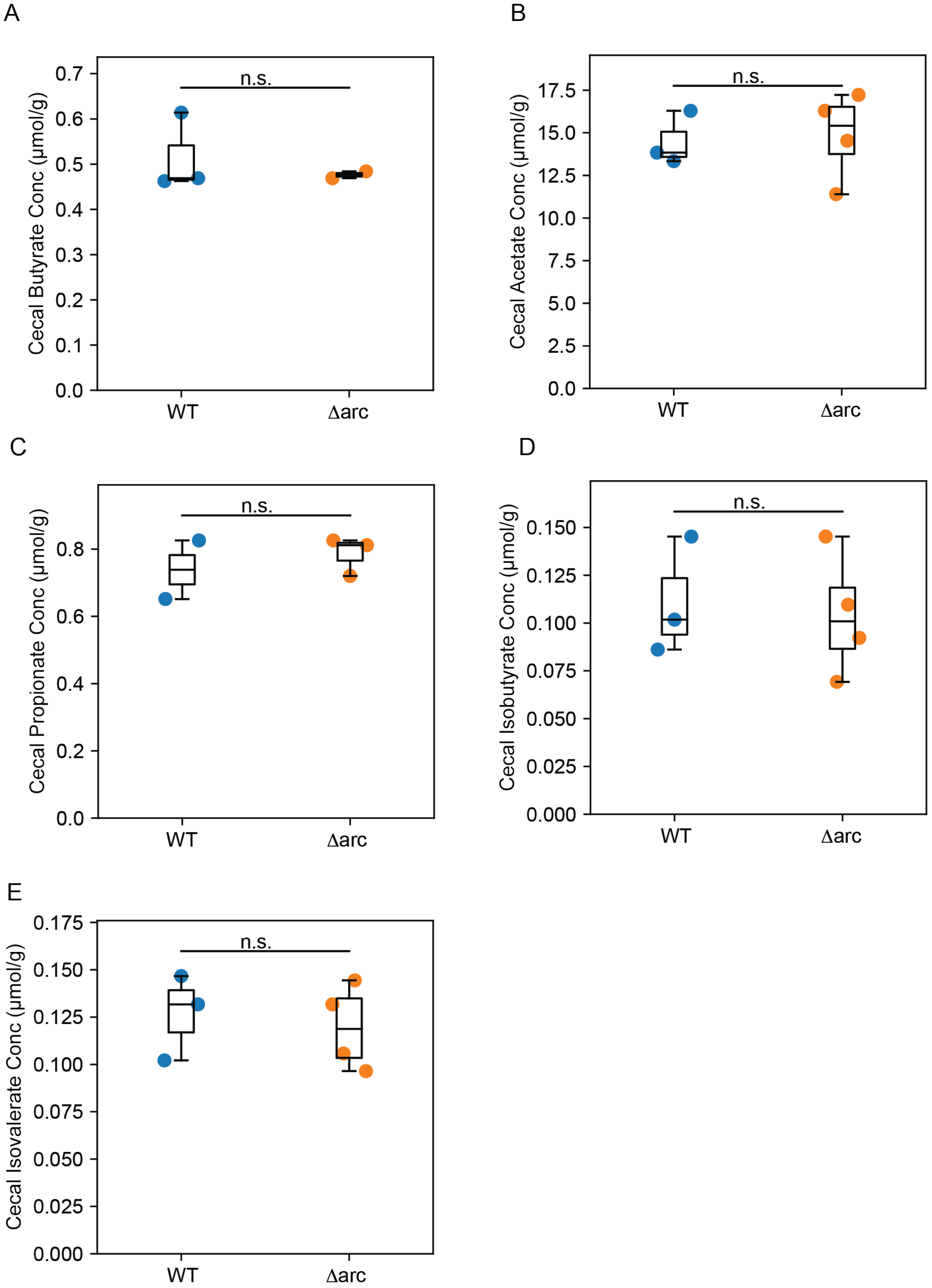


**Supplementary Figure 14. Metabolite measurements on cecal contents from mice.** Box plot of cecal butyrate (A), acetate (B), propionate (C), isobutyrate (D), and isovalerate (E) concentrations measured by gas chromatography. Cecal contents were harvested two weeks after introduction of EcN. Colors represent groups of germ-free mice colonized with community and various strains of EcN (WT and Δarc). The text n.s. represent no statistical significance according to an unpaired t-test between WT condition and Δarc condition.
